# Chondrocyte-Specific Knockout of *Piezo1* and *Piezo2* Protects Against Post-Traumatic Osteoarthritis Structural Damage and Pain in Mice

**DOI:** 10.1101/2025.05.22.655585

**Authors:** Erica V. Ely, Kristin L. Lenz, Sophie G. Paradi, Seth Ack, Abraham Behrmann, Sarah Dunivan, Lauryn Braxton, Wolfgang Liedtke, Yong Chen, Kelsey H. Collins, Farshid Guilak

**Author notes:** **Corresponding Author:** Farshid Guilak Department of Orthopaedic Surgery Washington University in St. Louis St. Louis, MO 63110.

## Abstract

**Background:** Osteoarthritis (OA) is a debilitating joint disease characterized by cartilage degeneration, synovial inflammation, and bone remodeling, with limited therapeutic options targeting the underlying pathophysiology. Mechanosensitive ion channels Piezo1 and Piezo2 play crucial roles in chondrocyte responses to mechanical stress, mediating mechanotransduction pathways that influence chondrocyte survival, matrix production, and inflammatory signaling, but their distinct contributions to OA pathogenesis remain unclear.

**Methods:** Using inducible, chondrocyte-specific Aggrecan-Cre (*Acan*) mice, we investigated *Piezo1*, *Piezo2*, and combined *Piezo1*/*2* conditional knockouts (cKOs) using the destabilization of the medial meniscus (DMM) model of post-traumatic OA in male and female mice. Pain and behavioral assessments were conducted at four time points to evaluate OA progression, while cartilage damage, bone remodeling, and synovial inflammation were assessed at the final endpoint of 28 weeks. Statistical analyses included one-way and two-way ANOVA with Tukey’s multiple comparisons test.

**Results:** *Piezo1* cKO delayed pain onset but ultimately exacerbated cartilage degradation and synovitis, emphasizing its dual role in protective and pathogenic mechanotransduction. While the *Piezo2* cKO reduced pain and preserved activity, it failed to protect cartilage. Notably, *Piezo1/2* cKO provided the greatest protection against cartilage degeneration, synovitis, and pain. Micro-computed tomography analyses revealed that *Piezo2* is critical for maintaining trabecular bone integrity, with a *Piezo2* cKO leading to decreased bone volume, thickness, and density, independent of injury. *Piezo2* cKO also reduced normal meniscal ossification that occurs with age in mice. In contrast, a *Piezo1/2* cKO normalized most bone remodeling parameters observed in *Piezo2* cKO mice but did not restore medial tibial plateau thickness, highlighting *Piezo2*’s essential role in bone structure.

**Conclusions:** These findings demonstrate the overlapping and compensatory roles of *Piezo1* and *Piezo2* in OA pathogenesis. Dual inhibition of *Piezo1 and Piezo2* may offer a novel, effective therapeutic strategy targeting both structural and symptomatic aspects of the disease.

## Introduction

Osteoarthritis (OA) is a painful and debilitating disease of the synovial joints that affects over 500 million people worldwide.^1^ It is characterized by degenerative changes in the morphology, composition, and mechanical properties of articular cartilage, as well as inflammation of the synovial fluid and alterations in bone structure.^2^ Pain and reduced activity levels are also common characteristics of OA, further contributing to decreased quality of life.^2,3^ Mechanical loading has a multi-faceted influence in joint health, as physiological loading is essential for maintaining cartilage homeostasis, while injurious or excessive loading – caused by joint instability, traumatic injury, or obesity – can disrupt cartilage integrity and accelerate OA progression.^4-7^ Understanding how mechanical forces are translated into biological signals within cartilage and surrounding tissues is essential for identifying the mechanobiological drivers of cartilage health, as well as OA, to allow for the development of targeted therapies to mitigate pain and tissue degeneration.

Chondrocytes, the primary cell type in cartilage, maintain homeostasis by sensing mechanical loads and translating these signals into intracellular responses through mechanosensitive ion channels.^8,9^ Mechanically activated cation channels such as Piezo1 and Piezo2 play central roles in this process, with calcium ion (Ca^²⁺^) signaling acting as a critical messenger that regulates downstream pathways influencing chondrocyte function and viability.^10,11^ Under physiological loading, these processes support cartilage health;^12^ however, high mechanical loading activates Piezo channels, leading to chondrocyte injury and apoptosis.^6,13-18^ In vitro studies have shown that inhibiting Piezo channels can protect chondrocytes from mechanical injury, suggesting their therapeutic potential.^11,14,19^ Piezo1 is primarily involved in mechanically induced chondrocyte cell death, potentially contributing to long-term cartilage degeneration,^16,18^ while Piezo2 in nociceptors mediates mechanical sensitization and pain responses^15,20,21^. Notably, *Piezo2* knockout reduces hyperalgesia and allodynia, highlighting its role in pain modulation.^13,22^ The synergistic function of *Piezo1* and *Piezo2* confers high-strain mechanosensitivity to cartilage, making these channels critical targets for understanding the mechanobiological drivers of osteoarthritis and for developing novel therapeutic strategies to mitigate disease progression.^18,19^

Despite significant advances, our understanding of the distinct contributions of *Piezo1* and *Piezo2* to OA pathogenesis remains incomplete, contributing to the lack of disease-modifying OA drugs.^2,17^ Prior in vivo studies have highlighted the complex roles of Piezo channels in OA. For example, a constitutive *Gdf5*-specific knockout of *Piezo1* and *Piezo2* resulted in moderate to severe OA and failed to protect joint integrity following destabilization of the medial meniscus (DMM) surgery.^23^ Conversely, intra-articular injection of GsMTx4, a Piezo channel inhibitor, ameliorated OA progression in a rat model following an anterior cruciate ligament transection (ACLT).^24^ Additionally, a constitutive *Col2a1*-specific knockout of *Piezo1*, but not *Piezo2*, in chondrocytes significantly attenuated cartilage degradation and inflammation after ACLT.^25^ An inducible *Acan*-specific knockout of *Piezo1* demonstrated protection from OA progression following DMM surgery, further supporting the potential therapeutic targeting of these channels.^26^ However, most studies have primarily focused on cartilage degradation, often neglecting the interconnected nature of physical joint damage — including synovitis, bone remodeling, and cartilage loss — and pain and behavioral outcomes. This limits our understanding of how the Piezo ion channels contribute to the broader pathophysiology of OA, impeding the development of holistic therapeutic strategies. To bridge this gap, further investigation is required to untangle the complex and sometimes contradictory roles of Piezo channels, enabling the design of targeted therapies that address both structural damage and symptomatic relief of OA.

The goal of this study was to address this knowledge gap by using an inducible, chondrocyte-specific, aggrecan-cre (*Acan)* driver to knock out *Piezo1*, *Piezo2*, or both channels simultaneously in a surgical model of post-traumatic OA. To comprehensively investigate the role of the Piezo ion channels in OA, we combine longitudinal assessments of pain and voluntary activity with histological evaluations of cartilage degeneration and joint inflammation, as well as structural analyses of subchondral bone changes. By examining both male and female mice, this study provides a multifaceted approach to understanding the contributions of *Piezo1* and *Piezo2* across different aspects of OA pathology. We hypothesized that cartilage-specific deletion of *Piezo1* and *Piezo2* will attenuate the progression of joint degeneration and pain, offering valuable insights into the development of novel disease-modifying OA therapies.

## Results

### qPCR Validation of *Piezo1* and *Piezo2* Knockout

To validate the efficiency of aggrecan-specific *Piezo1* and *Piezo2* knockout in cartilage, we conducted qPCR analysis of isolated hip cartilage from animals across the four genotypes: floxed control, *Piezo*1 conditional knockout (P1 cKO), *Piezo2* conditional knockout (P2 cKO), and *Piezo1/2* double conditional knockout (P1/2 cKO). For *Piezo1* expression, we observed significant differences between groups (n = 6-10/group, P < 0.001) (**Figure 1A**). P1 cKO mice exhibited a marked reduction in *Piezo1* expression compared to floxed controls (mean difference = 0.95, P < 0.001). Additionally, P1/2 cKO mice trended towards lower *Piezo*1 expression relative to floxed controls (mean difference = 0.63, P = 0.065). In contrast, *Piezo2* cKO mice demonstrated elevated *Piezo1* expression compared to *Piezo1/2* cKO (mean difference = 3.6, P < 0.001), *Piezo1* cKO (mean difference = 3.9, P < 0.001), and the floxed control group (mean difference = 2.9, P < 0.001) suggesting possible compensatory mechanisms. These results confirm that we successfully knocked out *Piezo1* in both *Piezo1* cKO and *Piezo1/2* cKO mice. For *Piezo2* expression, we also identified significant changes between groups (n = 6-10/group, P < 0.001) (**Figure 1B**). *Piezo2* expression was reduced in *Piezo2* cKO and *Piezo1/2* cKO mice versus both the floxed control and *Piezo1* cKO groups. Specifically, *Piezo2* cKO mice had the lowest *Piezo2* expression, significantly lower than the floxed control group (mean difference = 0.66, P < 0.001). Similarly, the *Piezo1/2* cKO group also showed reduced expression compared to floxed controls (mean difference = 0.42, P < 0.001). *Piezo1* cKO mice had comparable *Piezo2* expression to floxed controls, indicating no impact on *Piezo2* expression by *Piezo1* deletion. RT-qPCR Cycle threshold values show the average expression of *r18s* (mean = 16.64), *Piezo1* (mean = 22.94) and *Piezo2* (mean = 28.02) in floxed control murine hip caps (**Figure 1C)**. Overall, qPCR results validate the successful, specific knockouts of *Piezo1* and *Piezo2* in chondrocytes, confirming the efficiency of the genetic models used in this study.

**Figure 1.**
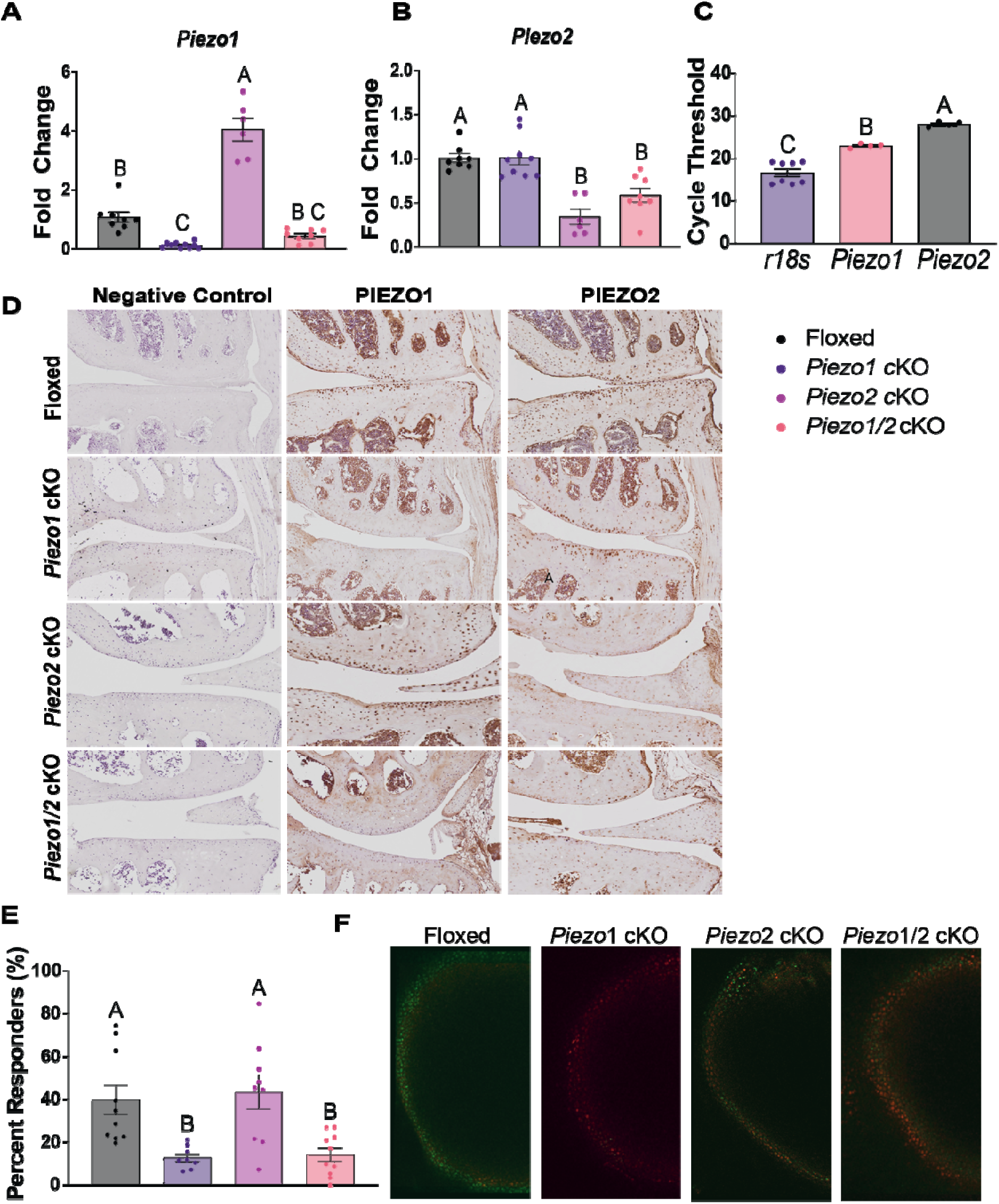
Validation of Piezo1 and Piezo2 Knockout and Functional Assessment. (A-C) Gene Expression Analysis: Quantitative PCR (qPCR) was performed on femoral condyle tissues from floxed (black), *Piezo1* cKO (P1 cKO-purple), *Piezo2* cKO (P2 cKO – pink), and Piezo1/2 cKO (P1/2 cKO – orange) mice at 28 weeks of age (n = 6-10/genotype). (A) *Piezo1* expression was significantly reduced in *Piezo1* cKO and *Piezo1/2* cKO mice while *Piezo2* cKO mice showed elevated Piezo1 expression. (B) *Piezo2* expression was significantly decreased in *Piezo2* cKO and *Piezo1/2* cKO groups compared to controls and *Piezo1* KO. Different letters represent statistical significance (one-way ANOVA with Tukey’s test, p < 0.05). (C) RT-qPCR cycle thresholds are shown for floxed control samples for *r18s* (housekeeping), *Piezo1*, and *Piezo2 (*n = 4-8/group). (D) Immunohistochemistry: Immunohistochemical staining of PIEZO1 and PIEZO2 was performed in the femoral condyle cartilage of each genotype. In floxed control mice, both PIEZO1 and PIEZO2 were expressed, while specific knockouts showed respective reductions in expression, represented by brown DAB staining. (E) Calcium Imaging Analysis: Functional knockout validation for *Piezo1* was assessed by quantifying calcium response following Yoda1 activation in chondrocytes. *Piezo1* cKO and *Piezo1/2* cKO mice demonstrated significantly lower responder percentages compared to floxed controls, indicating effective Piezo1 functional knockout. Different letters represent statistical significance (one-way ANOVA, p < 0.05). (F) Representative confocal images of calcium signaling in chondrocytes for floxed, *Piezo1* KO, *Piezo2* KO, and *Piezo1/2* cKO genotypes after Yoda1 perfusion. Fluo-4 (green) indicates calcium response. Scale Bar = 100μm.

To investigate potential off-target effects of the aggrecan-Cre driver in non-cartilage tissues, we evaluated *Piezo1* and *Piezo2* expression in the brain and lungs by qPCR. We observed a significant decrease in *Piezo2* expression in the lungs of *Piezo2* cKO mice, and a significant increase in *Piezo2* expression in the brains of *Piezo1* cKO mice (**Supplemental Figure 8A–D**), indicating some Cre activity outside of cartilage.

### Validation of *Piezo1* and *Piezo2* Knockout in Aggrecan-Expressing Tissues

We performed immunohistochemical staining using DAB (3,3’-diaminobenzidine) counterstain to assess PIEZO1 and PIEZO2 expression in aggrecan-expressing tissues, specifically targeting chondrocytes within the femoral condyle and tibial plateau (**Figure 1D**). In floxed control animals, chondrocytes within the femoral condyle and tibial plateau showed robust expression of both PIEZO1 and PIEZO2. In contrast, PIEZO1 staining was markedly reduced in *Piezo1* cKO mice, while PIEZO2 expression remained comparable to controls. Similarly, *Piezo2* cKO mice exhibited a significant reduction in PIEZO2 staining, with PIEZO1 expression remaining intact. Chondrocytes from *Piezo1/2* cKO mice displayed minimal to no staining for both PIEZO1 and PIEZO2, indicating effective knockout of these ion channels in chondrocytes. Negative control sections, processed without primary antibody, showed no background signal, confirming the specificity of the staining. These findings validate the successful and targeted knockout of PIEZO1 and/or PIEZO2 in chondrocytes.

To further examine whether PIEZO2 deletion could affect other cartilage-rich tissues involved in bone development, we assessed PIEZO2 expression in the femoral growth plates across all genotypes. As shown in **Supplemental Figure 8E**, PIEZO2 staining in the growth plate appeared comparable across floxed control, *Piezo1* cKO, *Piezo2* cKO, and *Piezo1/2* cKO mice, with no notable reduction in expression.

### Functional Validation of *Piezo* Knockout in Chondrocytes

We measured calcium signaling in response to Yoda1 treatment, a specific chemical activator of Piezo1, to confirm a functional knockout of the Piezo1 ion channels (n=8-10/genotype). *Piezo1* cKO mice showed significantly decreased calcium signaling versus floxed control mice (P < 0.004) and *Piezo2* cKO mice (P < 0.005) (**Figure 1E,F**). *Piezo1/2* cKO mice also displayed significantly lower calcium signaling compared to floxed control mice (P < 0.004) and *Piezo2* cKO mice (P < 0.005). In contrast, calcium signaling did not differ between *Piezo1* and *Piezo1/2* cKO mice (P < 0.998) or between floxed control and *Piezo2* cKO mice (P < 0.970). The *Piezo2* cKO group had a higher percent of responders, on average, than the floxed control group, potentially suggesting an increase in Piezo1 function following a *Piezo2* cKO. Overall, these results confirm a functional knockout of *Piezo1* in the *Piezo1* cKO and *Piezo1/2* cKO mice.

### Combined *Piezo1* and *Piezo2* Deletion Reduces OA Severity Following DMM Surgery

We evaluated Modified Mankin Scores in male mice 12 weeks after destabilization of the medial meniscus (DMM) surgery to the role of *Piezo1* and *Piezo2* on OA structural outcomes. In male mice, we observed a significant effect of surgery (P < 0.001), genotype (P = 0.010), and a significant interaction between genotype and surgery (P < 0.001) (n = 10-18/genotype). Modified Mankin scores were significantly higher in DMM limbs relative to contralateral control limbs across most genotypes, indicating an increase in cartilage damage after DMM surgery (**Figure 2A-B**). *Piezo1* cKO mice exhibited the highest Modified Mankin scores among all genotypes in the DMM limb (mean = 44 ± 4), exceeding their contralateral limb by a mean difference of 31 (P < 0.001), highlighting severe cartilage damage. Floxed control mice also demonstrated elevated scores in the DMM limbs (mean = 41 ± 2) compared to contralateral limbs (mean difference = 22, P < 0.001), reflecting substantial cartilage degradation. P2 cKO mice had increased scores in the DMM limb (mean = 40 ± 3) with a mean difference of 17 versus contralateral limb (P = 0.001), further confirming the significant impact of surgery on cartilage integrity. *Piezo1/2* cKO mice showed an intermediate response, with DMM limb scores (mean = 30 ± 2) exceeding the contralateral limbs by a mean difference of 10 (P = 0.008), suggesting a partial protective effect against cartilage damage with a combined knockout of *Piezo1* and *Piezo2*. *Piezo1/2* cKO DMM limbs had significantly lower scores relative to floxed controls (mean difference = 11, P = 0.002), *Piezo1* cKO (mean difference = 16, P < 0.001), and *Piezo2* cKO (mean difference = 10, P = 0.022) DMM limbs. In summary, the *Piezo1* cKO genotype appeared most susceptible to cartilage damage after DMM surgery, while the *Piezo1/2* cKO genotype exhibited a less pronounced increase in Modified Mankin scores, indicating protective effects of dual Piezo deletion in male mice.

**Figure 2.**
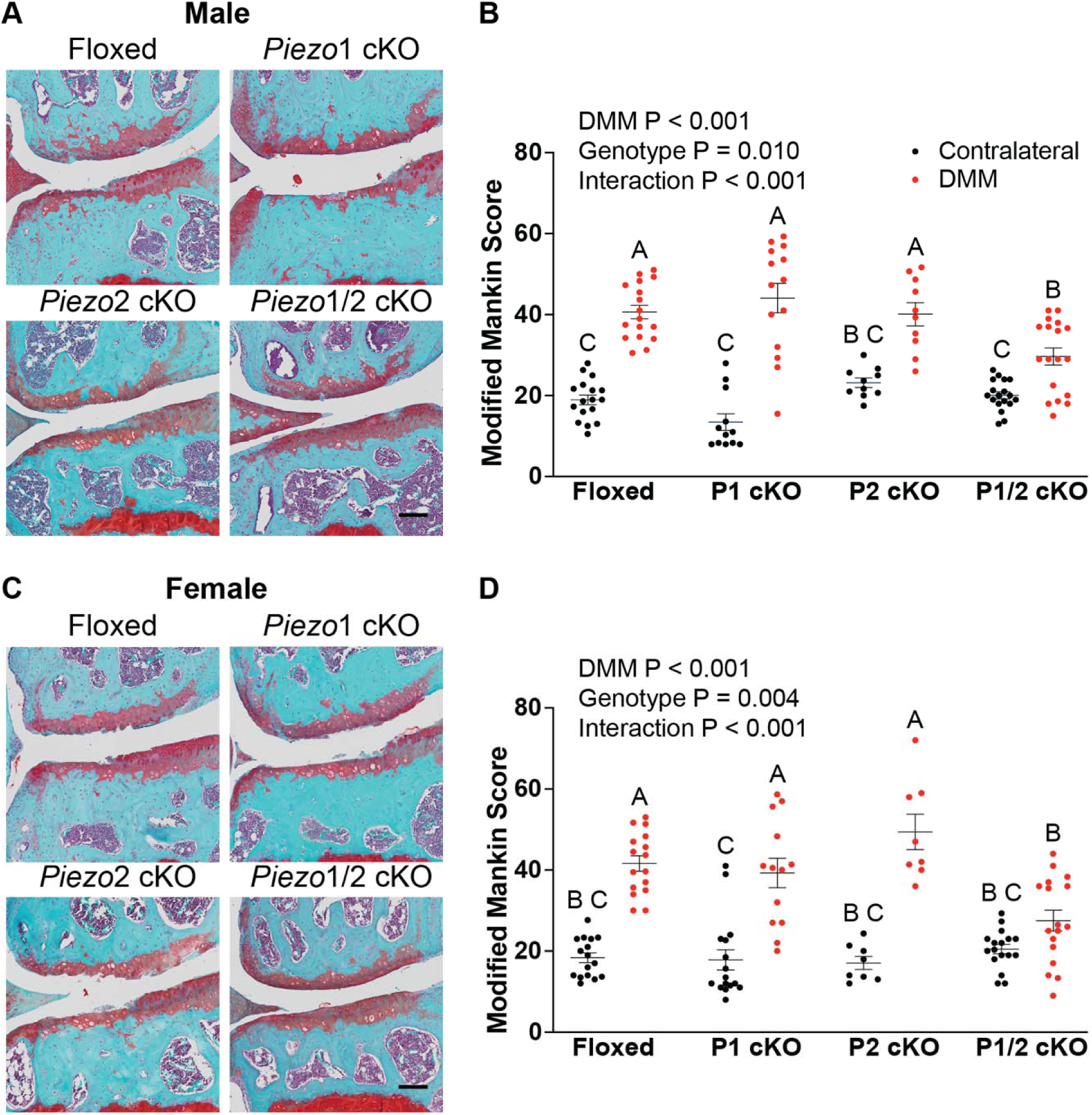
Histological Assessment of Cartilage Degradation Following DMM Surgery. (A, C) Representative images of coronal sections of the medial tibial plateau stained with Safranin-O/Fast from the knee joints of floxed, *Piezo1* KO, *Piezo2* KO, and *Piezo1/2* cKO male (A) and female (C) mice at 12 weeks post-DMM surgery. Red indicates cartilage, with more severe loss and lack of staining indicating greater cartilage degradation. (B, D) Quantification of Modified Mankin Scores for male (B) and female (D) mice. Red circles indicate scores for the DMM limb, while black circles represent the contralateral control limb. Data are presented as means ± SEM (n = 8-17/genotype). Different letters indicate statistically significant differences between groups based on a two-way ANOVA with Tukey’s multiple comparisons test (P < 0.05). For male mice, P1 cKO exhibited the highest Modified Mankin scores, while *Piezo1/2* cKO showed partial protection against cartilage damage compared to other groups. For female mice, *Piezo2 cKO* and *Piezo1* cKO exhibited severe cartilage damage, whereas *Piezo1/2* cKO demonstrated a less pronounced increase in Modified Mankin scores, suggesting a potential protective effect. Scale Bar = 100μm.

In female mice, we observed significant effects of surgery (P < 0.001), genotype (P = 0.004), and their interaction (P < 0.001) (n = 8-17/genotype). Modified Mankin scores were significantly higher in DMM limbs compared to contralateral control limbs across in all genotypes except for *Piezo1/2* cKO (**Figure 2C-D**). Specifically, P2 cKO mice had the highest Modified Mankin scores among all genotypes (mean = 49 ± 4) with a mean difference of 32 compared to their contralateral limbs (P < 0.001). *Piezo1/2* cKO mice DMM limb scores (mean = 39 ± 4) were significantly higher than their contralateral controls (mean difference = 21, P < 0.001), underscoring the impact of surgery on cartilage integrity. Floxed control mice similarly increased scores in the DMM limbs (mean = 42 ± 2) with a mean difference of 23 relative to contralateral limbs (P < 0.001). *Piezo1/2* cKO mice showed a mild response to surgery, with DMM limb scores (mean = 28 ± 3) not differing significantly from contralateral limbs (mean difference = 7, P = 0.299), suggesting a protective effect against OA. *Piezo1/2* cKO DMM limbs had significantly lower scores compared to floxed controls (mean difference = 14, P = 0.004), *Piezo1* cKO (mean difference = 12, P = 0.012), and *Piezo2* cKO (mean difference = 22, P < 0.001) DMM limbs. In summary, the *Piezo1* cKO and *Piezo2* cKO genotypes demonstrated the highest susceptibility to cartilage damage following DMM surgery, while the *Piezo1/2* cKO genotype showed reduced Modified Mankin scores, indicating a protective effect in female mice.

### *Piezo1/2* cKO Mice are Protected Against Increased Synovial Inflammation following DMM Surgery

We assessed synovitis scores using the Krenn criteria^27^ in male and female mice 12 weeks after destabilization of the medial meniscus (DMM) surgery. In male mice, there was a significant effect of surgery (P < 0.001) and the interaction between genotype and surgery (P = 0.014) (n = 9-20/genotype). DMM limbs displayed consistently higher synovitis scores than contralateral control limbs across all genotypes, indicating an increase in synovial inflammation after DMM surgery (**Figure 3A,C**). *Piezo2* cKO mice exhibited the highest synovitis scores in DMM limbs (mean = 6 ± 1), while contralateral limbs of *Piezo2* cKO mice had the lowest scores (mean = 2 ± 0.5). Floxed control mice demonstrated elevated synovitis scores in DMM limbs, with a mean difference of 2 compared to contralateral limbs (P = 0.029). *Piezo1* cKO mice showed increased synovitis in the DMM limb (mean = 5 ± 1), with a mean difference of 2 relative to the contralateral limb, though this difference was not statistically significant (P = 0.116). *Piezo1/2* cKO mice presented an intermediate response, with DMM synovitis scores (mean = 4 ± 0.5) slightly higher than the contralateral limbs (mean difference = 1, P = 0.937), but this change was also not significant.

**Figure 3:**
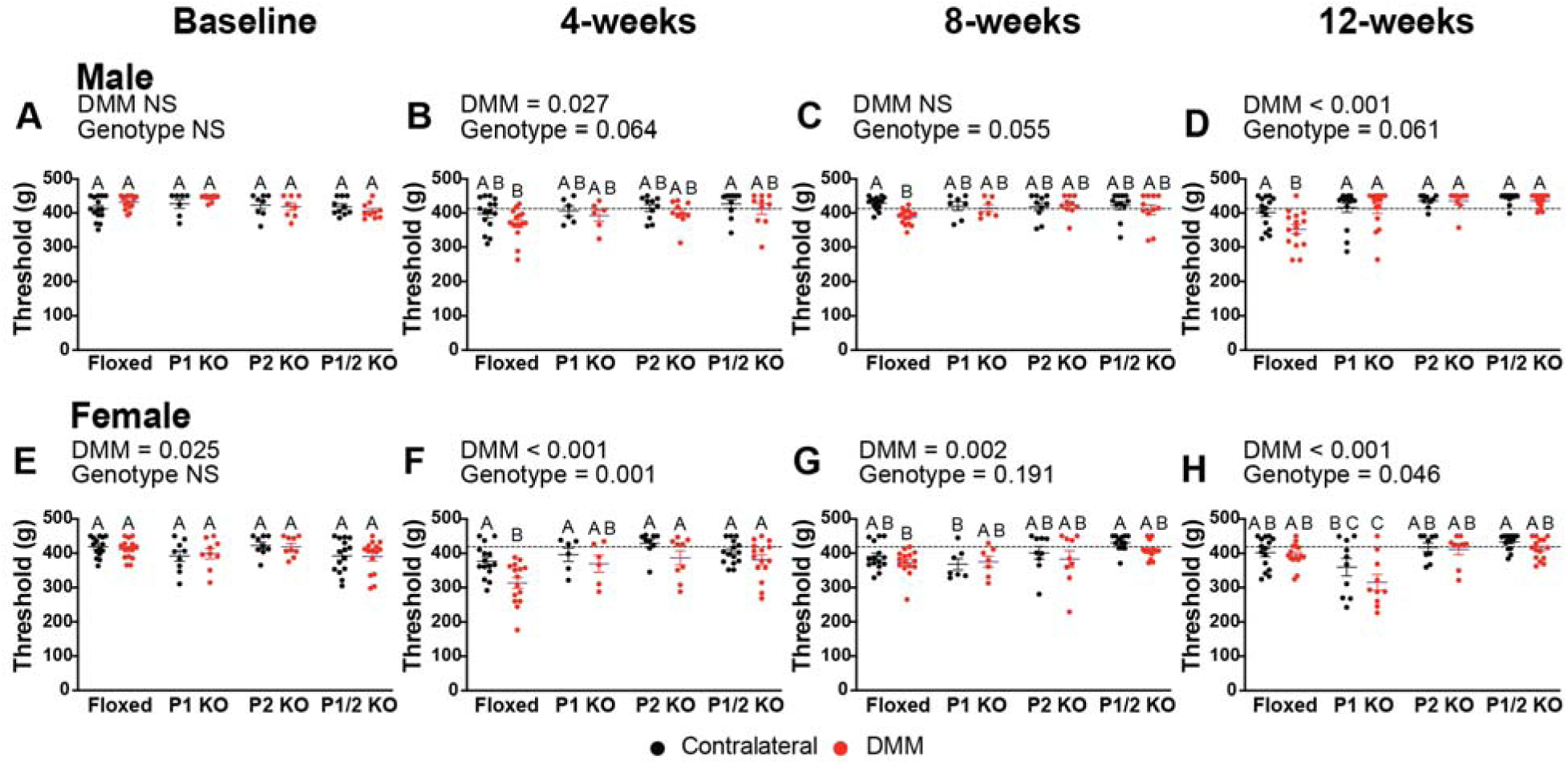
Osteophyte Severity and Synovitis Scores in Male and Female Mice 12 Weeks Post-DMM Surgery. (A-H) Representative H&E (A, C, E, G) and Safranin-O/Fast Green (B, D, F, H) stained sections of the medial tibial plateau regions are shown for floxed, *Piezo1* cKO, *Piezo2* cKO, and *Piezo1/2* cKO mice, highlighting synovitis severity and osteophyte formation. (C, G) Synovitis scores were quantified using the Krenn criteria (n = 9-20/genotype). In male mice (A,C), synovitis scores were significantly higher in DMM limbs compared to contralateral limbs in *Piezo2* cKO and floxed genotypes, while *Piezo1* cKO and *Piezo1/2* cKO exhibited intermediate responses. In female mice (E,G), synovitis scores were higher in DMM limbs across all genotypes, but *Piezo1/2* cKO did not show a significant difference compared to contralateral limbs.(D, H) Osteophyte numbers were quantified from Safranin-O/Fast Green stained sections (n = 7-15/genotype). In male mice (D), osteophyte numbers were significantly increased in DMM limbs of P1 cKO compared to contralateral limbs, while *Piezo1/2* cKO showed intermediate osteophyte numbers. No significant differences were observed in the other genotypes. In female mice (H), osteophyte numbers were higher in DMM limbs across all genotypes, but no significant differences were detected between groups or compared to contralateral limbs. Data are presented as mean ± SEM, with red dots representing DMM limbs and black dots representing contralateral limbs. Different letters indicate statistically significant differences (P < 0.05) based on two-way ANOVA and Tukey’s multiple comparisons test. Scale Bar = 200μm.

In female mice, a two-way ANOVA identified a significant effect of surgery (P < 0.001), but neither genotype alone (P = 0.056) nor the interaction between genotype and surgery (P = 0.123) reached statistical significance (n = 10-19/genotype). Synovitis scores increased in DMM limbs compared to contralateral control limbs in floxed control (mean difference = 3, P < 0.001), *Piezo1* cKO (mean difference = 3, P = 0.001), and *Piezo2* cKO mice (mean difference = 4, P < 0.001), reflecting heightened synovial inflammation after DMM surgery (**Figure 3E,G**). In contrast, *Piezo1/2* cKO mice did not exhibit a significant difference in synovitis scores between DMM and the contralateral limbs (mean difference = 2, P = 0.175). Among DMM limbs, *Piezo1* cKO mice showed the highest synovitis scores, indicating the most severe synovial inflammation. These findings suggest that the combined *Piezo1/2* knockout reduces synovitis severity in female mice, leading to less pronounced inflammation.

### Piezo Knockout Reduces Osteophyte Formation Post-DMM Surgery

We evaluated osteophyte formation in male mice following destabilization of the medial meniscus (DMM) surgery. In male mice, a two-way ANOVA identified significant effects of surgery (P < 0.001), genotype (P = 0.029), and their interaction (P = 0.036) (n = 9-15/genotype), highlighting the influence of both factors on osteophyte formation. Among the genotypes, only *Piezo1* cKO mice showed a significant increase in osteophyte numbers in DMM limbs compared to contralateral limbs (mean difference = 1.8, P < 0.001) (**Figure 3B,D**). *Piezo2* cKO, *Piezo1/2* cKO, and floxed control mice did not exhibit significant differences between limbs. The mean differences in osteophyte scores between contralateral and DMM limbs were 0.33 for *Piezo2* cKO, 0.95 for *Piezo1/2* cKO, and 0.71 for floxed control mice. When comparing DMM limbs across genotypes, *Piezo1* cKO mice had the highest osteophyte formation (mean = 1.83 ± 0.5), significantly exceeding *Piezo2* cKO DMM limbs (mean difference = 1.5, P = 0.005). *Piezo2* cKO and floxed control mice displayed the lowest osteophyte numbers in DMM limbs, while *Piezo1/2* cKO mice exhibited intermediate levels, which did not differ significantly from *Piezo2* cKO or floxed control mice. These findings suggest that a *Piezo1* cKO leads to the most pronounced osteophyte formation after joint destabilization and *Piezo1* and *Piezo2* play distinct, and possibly opposing, roles in osteophyte development.

In female mice, a two-way ANOVA revealed a significant effect of surgery (P = 0.023), indicating that DMM surgery increased osteophyte formation relative to contralateral control limbs (n = 7-15/genotype). However, genotype alone (P = 0.435) and its interaction with surgery (P = 0.889) did not reach statistical significance. Osteophytes remained consistently higher in DMM limbs compared to contralateral limbs across all genotypes (**Figure 3F,H**). The mean differences in osteophyte numbers between DMM and contralateral limbs were 0.42 for floxed control mice, 0.56 for *Piezo1* cKO mice, 0.14 for *Piezo2* cKO mice, and 0.43 for *Piezo1/2* cKO mice. When comparing DMM limbs across genotypes, no significant differences emerged, indicating similar levels of osteophyte formation across the board. These results suggest that, in female mice, osteophyte development after joint destabilization is influenced by surgery but not significantly by genotype.

### *Piezo2* Knockout Diminishes Bone Architecture and Density Post-DMM Surgery

We examined bone volume/total volume (BV/TV) in the tibial medial plateau of male mice (n = 9–16/genotype) and found a significant effect of genotype (P < 0.001), but no significant effects of surgery or interaction, indicating genotype differences independent of surgical interaction (**Figure 4A**). Within genotypes, no differences emerged between contralateral and DMM limbs, confirming that surgery did not alter BV/TV. However, the *Piezo2* cKO group displayed significantly lower BV/TV than the contralateral limbs of floxed control (P = 0.002), *Piezo1* cKO (P < 0.001), and *Piezo1/2* cKO (P = 0.031) mice. Across DMM limbs, *Piezo2* cKO mice had lower BV/TV than other genotypes, including floxed control (P = 0.048), *Piezo1* cKO (P = 0.001), and *Piezo1/2* cKO (P = 0.034), suggesting that *Piezo2* knockout in cartilage disrupts bone density at the medial tibial plateau.

**Figure 4.**
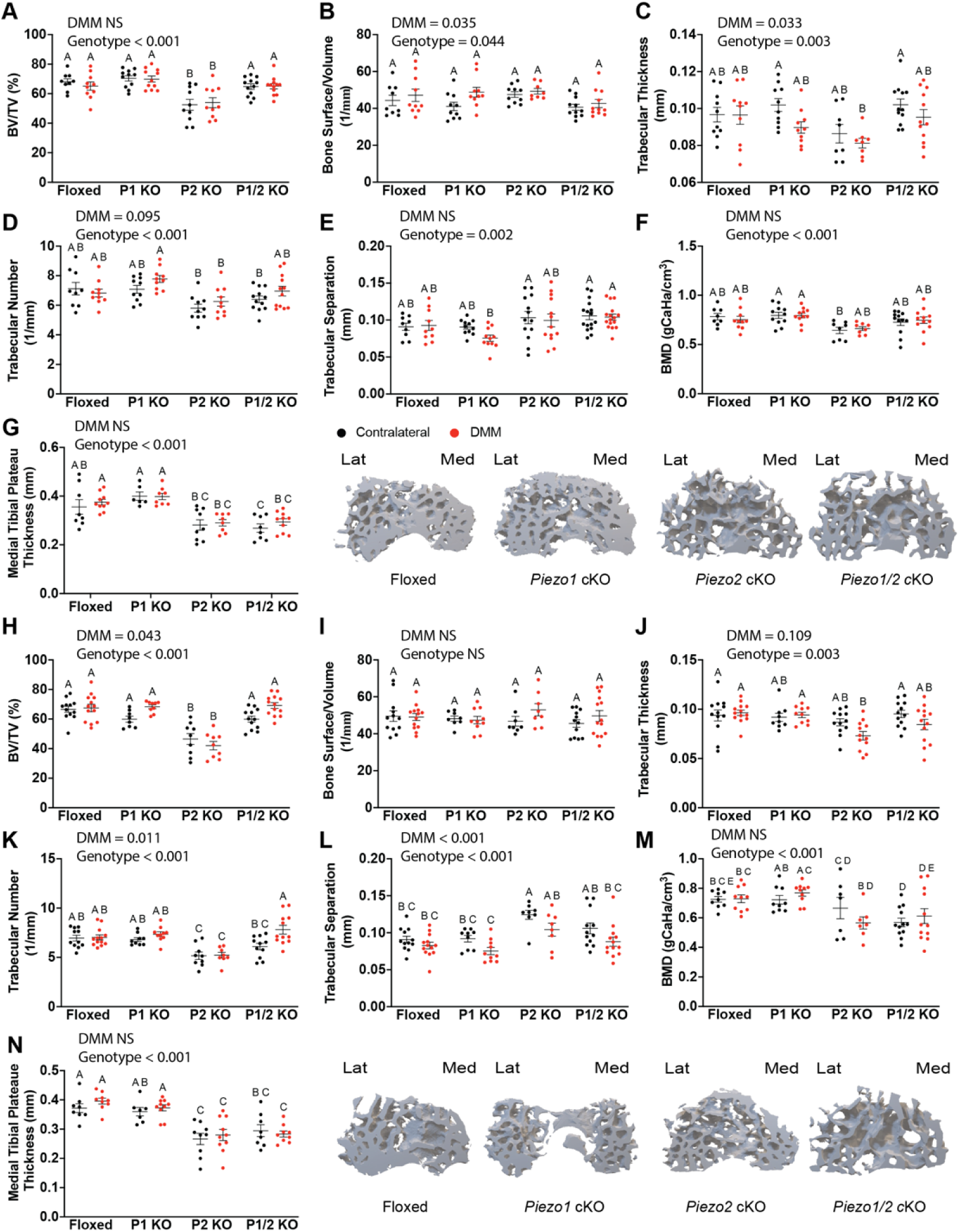
MicroCT Analysis of Bone Parameters in Male and Female Mice Following DMM Surgery. Bone morphometric outcomes were assessed in male (A-G) and female (H-N) Floxed, *Piezo1* cKO (P1 cKO), *Piezo2* cKO (P2 cKO), and *Piezo1/2* cKO (P1/2 cKO) mice at 12 weeks post-DMM. (A, H) Bone volume/total volume (BV/TV); (B, I) Bone surface/volume (BS/BV); (C, J) Trabecular thickness (Tb.Th); (D, K) Trabecular number (Tb.N); (E, L) Trabecular separation (Tb.Sp); (F, M) Bone mineral density (BMD); (G, N) Subchondral bone thickness (SBT) and representative 3D models of tibial plateau trabecular bone architecture. Data represent mean ± SEM (n=6-16/genotype/limb). Significant effects of genotype, surgery, and genotype-surgery interaction were tested using two-way ANOVA. Different letters indicate statistically significant differences (P < 0.05).

We assessed bone surface/volume (BS/BV) in male mice and found significant effects of genotype (P = 0.044) and surgery (P = 0.035) (**Figure 4B**). The lack of significant interaction (P = 0.557) indicated independent contributions of genotype and surgery. Contralateral and DMM limbs showed no differences within genotypes, suggesting that surgery itself did not alter BS/BV in the medial tibial plateau for any genotype.

For Trabecular thickness (Tb.Th), both genotype (P = 0.003) and surgery (P = 0.033), influenced outcomes, without significant interaction (P = 0.495) (**Figure 4C**). Surgery did not affect trabecular thickness within any genotype. However, *Piezo2* cKO mice exhibited lower trabecular thickness compared to the contralateral limbs of *Piezo1* cKO (P = 0.016) and *Piezo1/2* cKO (P = 0.001), suggesting *Piezo2* knockout uniquely alters trabecular architecture.

We analyzed trabecular number (Tb.N) in male mice and detected a significant effect of genotype (P = 0.001) but no effects of surgery (P = 0.095) or interaction (P = 0.360) (**Figure 4D**). Surgery did not alter Tb.N within genotypes. Across DMM limbs, *Piezo2* cKO mice had lower Tb.N than *Piezo1* cKO DMM (P = 0.012) and contralateral limbs (P = 0.022). Similarly, *Piezo2* cKO contralateral limbs displayed fewer trabeculae than *Piezo1* cKO DMM limbs (P = 0.003). These findings highlight *Piezo2* knockout as a significant factor in Tb.N reduction, independent of surgery.

Trabecular separation (Tb.Sp) results revealed significant effects of genotype (P = 0.0019) but no effects of surgery (P = 0.282) or interaction (P = 0.588) (**Figure 4E**). Contralateral and DMM limbs did not differ within genotypes. Among DMM limbs, *Piezo1* cKO mice showed lower trabecular separation compared to *Piezo2* cKO contralateral limbs (P = 0.039), *Piezo1/2* cKO contralateral limbs (P = 0.011), and *Piezo1/2* cKO DMM limbs (P = 0.020), emphasizing the distinct impact of *Piezo1* knockout on trabecular spacing.

Bone mineral density (BMD) results in male mice indicated a significant genotype effect (P < 0.001) but no effects of surgery (P = 0.917) or interaction (P = 0.849) (**Figure 4F**). Within genotypes, BMD remained consistent between contralateral and DMM limbs. The *Piezo2* cKO contralateral group displayed reduced BMD relative to *Piezo1* cKO contralateral (P = 0.026) and DMM limbs (P = 0.028). These findings highlight *Piezo2* knockout as a driver of reduced bone density, unaffected by surgery.

We measured medial tibial plateau thickness and found a significant genotype effect (P < 0.001), without effects of surgery (P = 0.309) or the interaction (P = 0.890) (**Figure 4G**). Thickness did not differ between contralateral and DMM limbs within genotypes. However, *Piezo1* cKO mice demonstrated significantly greater thickness than the *Piezo2* cKO contralateral (P = 0.001) and DMM limbs (P = 0.001). Floxed control DMM limbs showed greater thickness than *Piezo2* cKO contralateral (P = 0.005) and DMM (P = 0.020) limbs. The *Piezo1* cKO groups also had significantly greater thickness compared to the *Piezo1/2* cKO contralateral and DMM groups (P < 0.010 in all comparisons). These results suggest *Piezo2* knockout reduces tibial thickness, while *Piezo1* knockout may enhance or preserve it.

Given the striking bone changes observed in *Piezo2* cKO mice, we next quantified the ossified fraction of the menisci in the DMM limbs of floxed control, *Piezo1* cKO, *Piezo2* cKO, and *Piezo1/2* cKO mice. Our analysis revealed that *Piezo2* cKO male mice exhibited approximately 50% lower tissue volume (**Supplemental Figure 1A-C**) and bone volume (**Supplemental Figure 1D-F**) compared to floxed controls in the medial meniscus (P = 0.190, 0.191 respectively) and total meniscus (P = 0.058, 0.054 respectively). Additionally, *Piezo2* cKO mice had higher bone surface-to-volume ratio in the medial (P = 0.057) and total meniscus (P < 0.045) indicating thin bones and high trabecular numbers. *Piezo2* cKO mice also had significantly higher bone surface density compared to floxed controls in the medial (P < 0.019) and total (P < 0.011) meniscus indicating thin trabeculae and high porosity, emphasizing the distinct role of *Piezo2* in regulating bone remodeling.

We analyzed bone volume/total volume (BV/TV) in female mice (n = 9–13/genotype) and identified significant effects of genotype (P < 0.001), surgery (P = 0.043), and their interaction (P = 0.017) (**Figure 4H**). These results indicate that both genotype and surgical intervention influence BV/TV. The *Piezo2* cKO group exhibited significantly lower BV/TV than the contralateral limbs of floxed control (P < 0.001), *Piezo1* cKO (P = 0.014), and *Piezo1/2* cKO (P = 0.004) groups. Among DMM limbs, the *Piezo2* cKO group showed significantly reduced BV/TV compared to the other genotypes. Specifically, the *Piezo2* cKO group showed significantly lower BV/TV relative to floxed control (P < 0.001), *Piezo1* cKO (P = 0.002), and *Piezo1/2* cKO (P < 0.001) mice, suggesting that *Piezo2* knockout alters bone structure at the medial tibial plateau.

Bone surface/volume (BS/BV) in female mice revealed no significant effects of genotype (P = 0.772), surgery (P = 0.225), or interaction (P = 0.468) (**Figure 4I**). Neither genotype nor surgical intervention affected BS/BV, as all groups displayed comparable values between contralateral and DMM limbs, indicating no substantial alterations in this parameter.

We examined trabecular thickness (Tb.Th) and found a significant effect of genotype (P = 0.003) but no significant effects of surgery (P = 0.100) or interaction (P = 0.112) (**Figure 4J**). Surgery did not alter Tb.Th within genotypes. However, the P2 cKO group demonstrated significantly reduced trabecular thickness compared to the DMM limb of the floxed controls (P = 0.016) and the *Piezo1* cKO mice (P = 0.020), highlighting the impact of a *Piezo2* knockout on trabecular architecture.

Trabecular number (Tb.N) results in female mice indicated a significant effect of genotype (P < 0.001), surgery (P = 0.011), and their interaction (P = 0.029) (**Figure 4K**). The *Piezo1/2* cKO group displayed increased Tb.N in DMM limbs compared to contralateral limbs (P = 0.004), suggesting that surgery influenced this parameter in *Piezo1/2* cKO mice. Other genotypes showed no significant differences between contralateral and DMM limbs. Across genotypes, both *Piezo2* cKO contralateral and DMM groups had reduced Tb.N compared to the *Piezo1/2* cKO DMM group (P < 0.001). Additionally, the *Piezo1* cKO DMM group exhibited higher Tb.N than both *Piezo2* cKO groups (P < 0.001), illustrating genotype-dependent effects on bone architecture.

We investigated trabecular separation (Tb.Sp) and observed significant effects of genotype (P < 0.001) and surgery (P = 0.004), but not the interaction (P = 0.733) (**Figure 4L**). *Piezo2* cKO mice showed elevated Tb.Sp compared to the contralateral limbs of the floxed control (P = 0.006) and *Piezo1* cKO (P = 0.010), as well as the DMM limbs of floxed control (P = 0.002) mice. *Piezo2* cKO contralateral limbs also differed significantly from the *Piezo1/2* cKO DMM limbs (P = 0.002), underscoring distinct changes in Tb.Sp caused by *Piezo2* knockout.

Bone mineral density (BMD) results in female mice (n = 8-12/genotype/limb) highlighted a significant genotype effect (P < 0.001), but no impact from surgery (P = 0.785) (**Figure 4M**). Pairwise comparisons showed reduced BMD in the *Piezo1/2* cKO contralateral group compared to the floxed control contralateral limbs (P = 0.024), floxed control DMMs (P = 0.004), *Piezo1* cKO contralateral limbs (P = 0.043), and *Piezo1* cKO DMMs (P = 0.006). Similarly, the *Piezo2* cKO contralateral group exhibited lower BMD than the *Piezo1* cKO contralateral (P = 0.001) and DMM (P = 0.001) groups, indicated that *Piezo2* knockout reduces BMD independent of surgery.

Medial tibial plateau thickness analysis revealed a significant effect of genotype (P < 0.001) but no effects of surgery (P = 0.370) or interaction (P = 0.708) (n = 7/10/genotype) (**Figure 4N**). *Piezo2* cKO contralateral and DMM limbs displayed reduced thickness compared to the contralateral limbs of floxed control (P = 0.003 and P = 0.002, respectively), floxed control DMM (P < 0.001 for both), *Piezo1* cKO contralateral (P = 0.003 and P = 0.01, respectively), and *Piezo1* cKO DMM (P < 0.001 for both). *Piezo1/2* cKO contralateral and DMM limbs showed lower thickness than floxed control (P = 0.027 and P = 0.004, respectively) and floxed control DMM (P = 0.006 and P < 0.001, respectively). These findings emphasize the distinct effects of *Piezo2* and *Piezo1/2* knockouts on tibial architecture, regardless of surgical intervention.

### Grimace Scores Highlight Genotype-Specific Pain and Recovery After DMM Surgery

We evaluated grimace scores in male mice at baseline and at 4, 8, and 12 weeks post-DMM surgery to evaluate pain levels across four groups: Floxed controls, *Piezo1* cKO, *Piezo2* cKO, and *Piezo1/2* cKO (n = 7–15 per group) (**Figure** 5A**, Supplemental Figure 2A-D)**. At baseline (before surgery, 0 weeks), a one-way ANOVA showed no significant differences in grimace scores among the groups (P = 0.508), suggesting that pain levels were comparable across the genotypes prior to surgery (**Supplemental Figure 2A**). At 4 weeks post-DMM, all groups displayed increased grimace scores relative to baseline, indicating elevated pain following surgery, although no significant differences were observed among groups (P = 0.487) (**Supplemental Figure 2B**). At 8 weeks post-DMM, grimace scores remained elevated but consistent across genotypes, as a one-way ANOVA detected no significant differences (P = 0.846) (**Supplemental Figure 2C**). By 12 weeks post-DMM, significant differences emerged among the groups (P = 0.002) (**Figure 5A**, **Supplemental Figure 2D**). Tukey’s multiple comparisons test identified higher grimace scores in *Piezo2* cKO mice compared to floxed controls (adjusted P = 0.024) and *Piezo1/2* cKO mice (adjusted P = 0.001). Although *Piezo1* cKO trended towards higher scores than *Piezo1/2* cKO mice, the difference did not reach statistical significance (adjusted P = 0.120). All groups maintained scores above baseline, while the *Piezo1/2* cKO mice demonstrated recovery to levels closest to pre-surgery values (mean = 1.68). These results suggest that *Piezo2* cKO mice experienced the most pain, whereas *Piezo1/2* cKO mice exhibited better recovery by 12 weeks post-DMM, although not significantly different than floxed controls.

**Figure 5.**
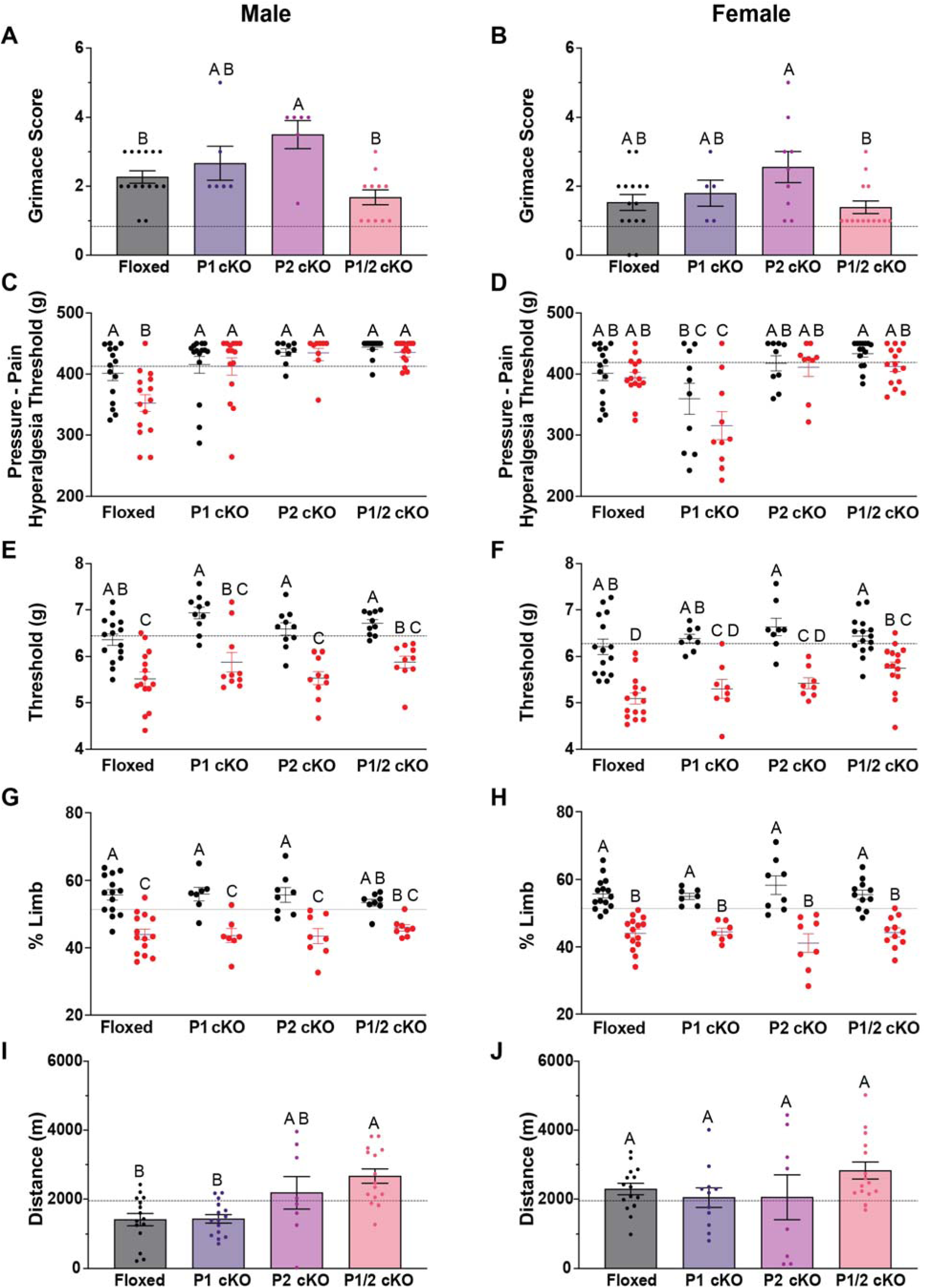
Pain and Activity Outcomes in Male and Female Mice 12-Weeks Post-DMM Surgery. Male (A) and female (B) Floxed, *Piezo1* cKO (P1 cKO - purple), *Piezo2* cKO (P2 cKO - pink), and *Piezo1/2* cKO (P1/2 cKO - orange) mice were assessed using the Mouse Grimace Scale (MGS) at 12 weeks post-DMM surgery. *Piezo2* cKO male mice exhibited higher pain scores compared to floxed and *Piezo1/2* cKO groups (P = 0.002) (A), while female *Piezo1/2* cKO mice had significantly lower pain scores compared to *Piezo2* cKO mice (P = 0.037) (B). Pressure-pain hyperalgesia thresholds were evaluated using the SMALGO in both DMM (red) and contralateral limbs (black) of male (C) and female (D) Floxed, *Piezo1* cKO (P1 KO), *Piezo2* cKO (P2 KO), and *Piezo1/2* cKO (P1/2 KO) mice 12 weeks post-DMM. A two-way ANOVA was used for statistical analysis. All *Piezo c*KO male groups have significantly higher DMM thresholds compared to floxed mice (C). In female mice, a *Piezo2 c*KO results in decreased thresholds compared to all groups (D). Tactile allodynia thresholds were evaluated using the Electronic Von Frey (EVF). All male groups had significantly lower DMM thresholds compared to contralateral limbs, with no significant differences across genotypes (E). In female mice, *Piezo1/2* cKO DMM thresholds were significantly higher compared to floxed mice, indicating a protective effect (F). Static weight bearing measurements were evaluated there were significant differences in weight distribution across most genotypes, except for *Piezo1/2 c*KO male mice (G), highlighting the persistent effects of DMM surgery in male and female mice (H). Voluntary wheel running distances (in meters) were assessed and there was a significant difference in distance run male groups at 12 weeks (P < 0.001) (I), where *Piezo1/2* cKO mice exhibited significantly greater activity compared to *Piezo1* cKO and floxed controls. For female mice, no significant differences were observed across genotypes at any timepoint, with all groups demonstrating comparable recovery in running distances post-DMM surgery (J). The dashed lines represent baseline, pre-surgical, pain and activity measures of floxed mice (A-J). Statistics were run using either a one-way (A, B, I, J) or two-way (C-H) ANOVA followed by Tukey’s multiple comparisons test to assess differences between genotypes and/or limbs (n=6-15/genotype/sex). Different letters above bars indicate significant differences between groups (P < 0.05). Error bars represent mean ± SEM.

We also assessed grimace scores in female mice at the same timepoints (n = 6– 16/genotype) (**Figure 5 B**, **Supplemental Figure 2E-H**). At baseline, significant differences appeared among the groups (P < 0.002) (**Supplemental Figure 2E**). Tukey’s multiple comparisons test revealed elevated scores in *Piezo2* cKO mice compared to floxed controls (adjusted P < 0.003), *Piezo1* cKO (adjusted P = 0.022), and *Piezo1/2* cKO (adjusted P < 0.003), indicating that *Piezo2* cKO alone is sufficient to elevate pain levels. At 4 weeks post-DMM, grimace scores increased across genotypes (P = 0.021) (**Supplemental Figure 2F**). Tukey’s test showed that *Piezo1* cKO mice had the lowest grimace scores among genotypes, with lower scores than *Piezo2* cKO mice (adjusted P = 0.041) and floxed controls (P = 0.056). All genotypes displayed scores above baseline, reflecting heightened pain following surgery. At 8 weeks post-DMM, significant differences persisted (P = 0.008) (**Supplemental Figure 2G**). Tukey’s test found higher scores in *Piezo2* cKO mice relative to floxed controls (adjusted P = 0.041) and *Piezo1* cKO mice (adjusted P = 0.011). *Piezo1* cKO mice showed scores near baseline, indicating lower pain levels, while other genotypes maintained scores above baseline. At 12 weeks post-DMM, significant differences remained (P = 0.037) (**Figure 5B, Supplemental Figure 2H**). Tukey’s test identified *Piezo2* cKO mice with higher scores than *Piezo1/2* cKO mice (adjusted P = 0.031). *Piezo1/2* cKO mice exhibited the lowest grimace scores (mean = 1.39), suggesting reduced pain levels, although not significantly different from floxed controls (mean = 1.53). Although all groups maintained scores above baseline at this timepoint, *Piezo1/2* cKO mice demonstrated the greatest protection against sustained pain.

### *Piezo2* and *Piezo1/2* Knockouts Mitigate Pressure-Pain Hyperalgesia

We assessed the pressure-pain hyperalgesia threshold at baseline across four groups of male mice: Floxed controls, *Piezo1* cKO, *Piezo2* cKO, and *Piezo1/2* cKO (n = 6–15 per genotype) (**Figure 5C, Supplemental Figure 3A-D**). At baseline, a two-way ANOVA revealed no significant differences among genotypes (P = 0.475), effects of surgery (P = 0.138) or interaction between genotype and surgery (P = 0.169) (**Supplemental Figure 3**A). At 4 weeks post-DMM, we reassessed pain thresholds and identified significant effects of surgery (P = 0.027), though genotype and interaction effects remained non-significant (interaction: P = 0.899; genotype: P = 0.064) (**Supplemental Figure 3B**). Post hoc Tukey’s comparisons showed no significant differences among groups, indicating stable pain thresholds relative to baseline values and consistent pain responses post-surgery. At 8 weeks post-DMM, thresholds remained stable across all groups. A two-way ANOVA did not detect significant effects of surgery (P = 0.61) or genotype (P = 0.055), and no interaction was observed (P = 0.078) (**Supplemental Figure 3C**). However, the floxed control DMM group exhibited a higher threshold than its contralateral limb (mean difference = 41.3 g, P = 0.015). Other groups maintained similar pain thresholds across limbs. These findings suggest that while the floxed controls showed slight changes in thresholds, other genotypes maintained consistency. By 12 weeks post-DMM, surgery significantly impacted thresholds (P < 0.001), but genotype effects (P = 0.061) and the interaction (P = 0.099) remained non-significant (**Figure 5C**, **Supplemental Figure 3D**). Tukey’s comparisons revealed that the floxed control DMM group had lower thresholds than the *Piezo1/2* cKO DMM (mean difference = 83.0 g, P < 0.001), the *Piezo2* cKO DMM (mean difference = 82.1 g, P = 0.002), and the *Piezo1* cKO DMM (mean difference = 59.9 g, P = 0.003) groups. *Piezo2* cKO and *Piezo1/2* cKO groups maintained thresholds above baseline, with the *Piezo1/2* cKO group consistently showing the highest thresholds across all timepoints. These results highlight persistent hypersensitivity and reduced thresholds in floxed controls while all Piezo cKO genotypes demonstrated greater resilience.

We evaluated female mice at the same timepoints (n = 6–15/genotype) (**Figure 5D, Supplemental Figure 3E-H**). At baseline, a two-way ANOVA showed no significant differences in thresholds among genotypes (P = 0.944) or interactions between surgery and genotype, though thresholds differed between DMM and contralateral limbs (P = 0.025) (**Supplemental Figure 3E**). At 4 weeks post-DMM, surgery (P = 0.003) and genotype (P = 0.001) significantly affected thresholds (**Supplemental Figure 3F**). The floxed control group displayed lower thresholds in the DMM limb relative to the contralateral limb (mean difference = 63.6 g, P = 0.021). The *Piezo1/2* cKO DMM group showed higher thresholds than floxed control DMM limbs (mean difference = 66.9 g, P = 0.012). All groups demonstrated thresholds below baseline, reflecting heightened sensitivity. At 8 weeks post-DMM, significant effects of surgery persisted (P = 0.002), though Tukey’s multiple comparison’s test did not reveal significant differences between limbs or genotypes (**Supplemental Figure 3G**). At 12 weeks post-DMM, both surgery (P < 0.001) and genotype (P = 0.046) significantly influenced thresholds (**Figure 5D, Supplemental Figure 3H**). Tukey’s test highlighted higher pain threshold in the floxed control DMM limbs compared to the *Piezo1* cKO DMM group (mean difference = 78.9 g, P = 0.002), indicating increased hypersensitivity in the *Piezo1* cKO group. *Piezo1* cKO DMM limbs displayed lower thresholds than *Piezo2* cKO DMM (mean difference = 95.7 g, P = 0.006) and *Piezo1/2* cKO DMM (mean difference = 96.9 g, P = < 0.001) limbs. *Piezo2* cKO and *Piezo1/2* cKO mice consistently exhibited the highest pain thresholds overall, closest to baseline, indicating reduced pain development. All groups showed reduced thresholds relative to baseline, with significant hypersensitivity most evident in *Piezo1* cKO mice.

### Genotype-Specific Tactile Allodynia Responses Illustrate *Piezo1/2* Combined Role in OA pain

We assessed mechanical (or tactile) pain thresholds across all groups of male and female mice: Floxed controls, *Piezo1* cKO, *Piezo2* cKO, and *Piezo1/2* cKO (n=7–15/genotype). At baseline, a two-way ANOVA detected no significant differences among genotypes (P = 0.179), no effects of surgery (P = 0.422), and no interaction between surgery and genotype (P = 0.819) (**Supplemental Figure 4A**). These results indicate uniform baseline pain sensitivity across genotypes before the DMM surgery. At 4 weeks post-DMM, thresholds were reevaluated and a two-way ANOVA revealed significant effects of surgery (P = 0.038) and genotype (P = 0.001) but no interaction (P = 0.547) (**Supplemental Figure 4B**). Post hoc Tukey’s multiple comparisons showed no significant changes between contralateral and DMM limbs within genotypes or among genotypes for DMM limbs, though floxed controls trended towards lower thresholds in DMM limbs (mean difference = 0.658, P = 0.103). These data suggest that at 4 weeks, surgery begins influencing thresholds, though significant differences are not yet evident. At 8 weeks post-DMM, a two-way ANOVA identified significant effects of genotype (P < 0.001), but neither surgery (P = 0.249) nor interactions between factors (P = 0.113) were significant (**Supplemental Figure 4C**). Post hoc Tukey’s tests revealed significantly lower thresholds in DMM limbs for the floxed control, *Piezo2* KO, and *Piezo1/2* cKO groups (floxed control mean difference = 0.789, P = 0.002, *Piezo2* cKO mean difference = 0.640, P = 0.046, *Piezo1/2* cKO mean difference = 1.19, P < 0.001). *Piezo1* cKO male mice did not exhibit significant differences between limbs, suggesting a distinct role for *Piezo1* in pain onset. All DMM groups showed thresholds below baseline levels, reflecting persistent pain. At 12 weeks post-DMM, thresholds varied significantly due to genotype (P < 0.001) and surgery (P = 0.005), though no interaction was observed (P = 0.769) (**Figure 5E**, **Supplemental Figure 4D**). Tukey’s test revealed lower thresholds in DMM limbs compared to contralateral limbs for all groups: Floxed controls (mean difference = 0.842, P = 0.002), *Piezo1* cKO (mean difference = 1.06, P = 0.001), *Piezo2* cKO (mean difference = 1.05, P = 0.001), *Piezo1/2* cKO (mean difference = 0.830, P = 0.006). Across genotypes, no significant differences emerged among DMM limbs. All DMM groups remained below baseline levels, confirming sustained pain post-surgery.

We assessed female mice at the same timepoints (n=8–15/genotype) (**Figure 5F, Supplemental Figure 4E-H**). At baseline, a two-way ANOVA found no significant differences among genotypes, surgery or interactions, indicating minimal pain before surgery (**Supplemental Figure 4E**). At 4 weeks post-DMM, a two-way ANOVA detected significant effects of genotype (P < 0.001) and surgery (P = 0.019) (**Supplemental Figure 4F**). Tukey’s multiple comparisons highlighted lower thresholds in the floxed control DMM limb relative to its contralateral limb (mean difference = 0.758, P = 0.021). Although other comparisons between limbs did not show significance, all DMM limb averages fell below baseline levels, indicating increased sensitivity. At 8 weeks post-DMM, surgery (P < 0.001) and genotype (P < 0.001) significantly influence thresholds (**Supplemental Figure 4G**). All groups displayed lower thresholds in DMM limbs than contralateral limbs: Floxed controls (mean difference = 0.853, P < 0.001), *Piezo1* cKO (mean difference = 0.871, P = 0.008), *Piezo2* cKO (mean difference = 0.728, P = 0.004), and *Piezo1/2* cKO (mean difference = 0.729, P = 0.004). Among DMM limbs, *Piezo2* cKO thresholds exceeded those of floxed controls (mean difference = 0.498, P = 0.049), suggesting some protection from pain in *Piezo2* cKO mice. All DMM groups remained below baseline levels. At 12 weeks post-DMM, surgery (P = 0.005) and genotype (P < 0.001) continued to influence thresholds (**Figure 5F, Supplemental Figure 4H**). All groups exhibited lower thresholds in DMM limbs than contralateral limbs: Floxed control (mean difference = 1.11, P < 0.001), *Piezo1* cKO (mean difference = 1.08, P < 0.001), *Piezo2* cKO (mean difference = 1.21, P < 0.001), *Piezo1/2* cKO (mean difference = 0.684, P = 0.006). Among DMM limbs, *Piezo1/2* cKO thresholds trended towards being higher than those of floxed control DMM limbs (mean difference = 0.658, P = 0.010), indicating better pain mitigation. While all groups showed thresholds below baseline values, the *Piezo1/2* cKO genotype demonstrated the best protection, underscoring the role of *Piezo1* and *Piezo2* in pain progression for female mice.

### Genotype-Dependent Static Weight Bearing Changes in Male and Female Mice Post-DMM Surgery

We evaluated static weight bearing in male mice from four groups: Floxed controls, *Piezo1* cKO, *Piezo2* cKO, and *Piezo1/2* cKO—at baseline (0 weeks) and 4, 8, and 12 weeks post-DMM surgery (n = 6–15/genotype) (**Figure 5G, Supplemental Figure 5A-D**). At baseline, a two-way ANOVA identified a significant effect of genotype on static weight bearing (P = 0.005), but no significant differences emerged between contralateral and DMM limbs within any genotype, indicating balanced weight distribution across all groups (**Supplemental Figure 5A**). These results confirm that genotypes showed no impaired load distribution prior to DMM surgery. At 4 weeks post DMM, static weight bearing decreased in the DMM limb across most groups. A two-way ANOVA revealed a significant effect of genotype (P < 0.001), suggesting that the differences observed were dependent on genotype rather than surgery alone (**Supplemental Figure 5B**). Within genotypes, floxed controls (mean difference = 9.68%, P < 0.0001), *Piezo1* cKO (mean difference = 17.14%, P < 0.001), and *Piezo1/2* cKO (mean difference = 13.59%, P < 0.001) mice exhibited significant reductions in DMM limb weight bearing, favoring the contralateral limb. For *Piezo2* cKO mice, the reduction did not reach significance (mean difference = 7.81%, P = 0.168). Additionally, all groups demonstrated DMM limb weight bearing below baseline levels observed in floxed controls, highlighting impaired load distribution following DMM surgery. Across genotypes, no significant differences were observed in DMM limb scores. At 8 weeks post DMM, a two-way ANOVA detected a significant interaction between genotype and limb (P = 0.048) and a significant effect of genotype (P < 0.001), suggesting the combined influence of genotype and limb on weight bearing (**Supplemental Figure 5C**). Within genotypes, floxed controls (mean difference = 16.23%, P < 0.0001), *Piezo1* cKO (mean difference = 19.65%, P < 0.001), *Piezo2* cKO (mean difference = 17.23%, P < 0.001), and *Piezo1/2* cKO (mean difference = 10.90%, P < 0.001) showed significant differences between contralateral and DMM limbs. DMM limb scores in all groups remained below the baseline load of floxed controls, emphasizing the impact of DMM surgery on load distribution. No significant differences emerged among genotypes for DMM limb scores. At 12 weeks post-DMM, prior to sacrifice, a two-way ANOVA revealed a significant effect of genotype (P < 0.0001), indicating genotype-dependent differences in static weight bearing (**Figure 5G, Supplemental Figure 5D**). Within genotypes, floxed controls (mean difference = 11.72%, P < 0.001), *Piezo1* cKO (mean difference = 12.27%, P = 0.002), and *Piezo2* cKO (mean difference = 12.20%, P < 0.001) mice exhibited significant reductions in DMM limb weight bearing compared to contralateral limbs. However, *Piezo1/2* cKO mice did not show significant differences between limbs (mean difference = 7.50%, P = 0.113). DMM limb scores for all genotypes remained below levels of floxed control mice, confirming impaired load distribution favoring the contralateral limb. These results indicate that both Piezo1 and Piezo2 ion channels play a role in maintaining normal load-bearing capacity post-DMM surgery in male mice.

Female mice displayed similar trends in static weight bearing across timepoints (n = 6– 15/genotype) (**Figure 5H**, **Supplemental Figure 5E-H**). At baseline, a two-way ANOVA detected significant genotype differences (P = 0.004), but contralateral and DMM limbs within each genotype maintained even weight distribution (**Supplemental Figure 5E**). These results confirm consistent static weight bearing across genotypes prior to joint injury. At 4-weeks post DMM, static weight bearing declined in DMM limbs across most groups. A two-way ANOVA revealed significant interactions between genotype and limb (P = 0.031) and significant effects of genotype (P < 0.001) (**Supplemental Figure 5F**). Within genotypes, floxed controls (mean difference = 17.72%, P < 0.001), *Piezo2* cKO (mean difference = 17.29%, P < 0.001), and *Piezo1/*2 cKO (mean difference = 17.64%, P < 0.001) showed significant differences between limbs. *Piezo1* cKO mice did not exhibit significant differences (mean difference = 7.20%, P = 0.313). DMM limb scores across all groups fell below baseline levels in floxed controls, indicating impaired load distribution. At 8-weeks post DMM, static weight bearing remained reduced. A two-way ANOVA identified a significant effect of genotype (P < 0.001) (**Supplemental Figure 5G**). Within genotypes, floxed controls (mean difference = 12.84%, P < 0.001), *Piezo1* cKO (mean difference = 11.44%, P = 0.038), *Piezo2* cKO (mean difference = 13.47%, P < 0.001), and *Piezo1/2* cKO mice (mean difference = 9.59%, P = 0.001) demonstrated significant differences between contralateral and DMM limbs. All DMM limb scores remained below baseline levels, reinforcing the impact of DMM surgery on load distribution. Comparisons across genotypes showed no significant differences in DMM limb scores. At 12-weeks post DMM, a two-way ANOVA indicated a significant genotype effect (P < 0.001) (**Figure 5H, Supplemental Figure 5H**). Within genotypes, floxed controls (mean difference = 11.67%, P < 0.001), *Piezo1* cKO (mean difference = 10.47%, P = 0.007), *Piezo2* cKO (mean difference = 17.18%, P < 0.001), and *Piezo1/2* cKO (mean difference = 11.18%, P < 0.001) mice exhibited significantly lower DMM limb scores than contralateral limbs. All DMM limb scores remained below baseline levels, confirming impaired load distribution post-surgery. No differences emerged among genotypes in DMM limb scores.

### Recovery of Voluntary Wheel Running Activity in *Piezo2* and *Piezo1/2* cKO Mice Post-DMM Surgery

We monitored voluntary wheel running distances in four groups of male mice: Floxed controls, *Piezo1* KO, *Piezo2* KO, and *Piezo1/2* cKO mice, across baseline (0 weeks), 4 weeks, 8 weeks, and 12 weeks post-DMM (n = 6–15/genotype) (**Figure 5I, Supplemental Figure 6A-D**). At baseline (0 weeks), a one-way ANOVA found no significant differences in distances run between groups (P = 0.321) (**Supplemental Figure 6A**), confirming comparable activity levels across genotypes before surgery. At 4 weeks post-DMM, a one-way ANOVA detected no significant differences in distances run between groups (P = 0.336) (**Supplemental Figure 6B**). While the floxed control and *Piezo1* cKO groups ran slightly less than the baseline average, *Piezo2* cKO and *Piezo1/2* cKO mice maintained distances above baseline. However, the lack of significance indicates minimal changes in activity levels following surgery. At 8 weeks post-DMM, a one-way ANOVA again indicated no significant differences between the groups (P = 0.523) (**Supplemental Figure 6C**). All genotypes ran on average, further than baseline levels, indicating little change on activity levels 8 weeks following DMM surgery. At 12 weeks post-DMM, a one-way ANOVA identified significant differences in distances run between groups (P < 0.001) (**Figure 5I, Supplemental Figure 6D**). Tukey’s multiple comparisons test showed that *Piezo1/2* cKO mice ran significantly farther than floxed controls (adjusted P < 0.001) and *Piezo1* cKO mice (adjusted P < 0.001). The distance run by *Piezo1* cKO mice and *Piezo2* cKO mice did not differ significantly (adjusted P = 0.153). Notably, *Piezo2* cKO and *Piezo1/2* cKO mice exceeded baseline control values, indicating a recovery to pre-surgery activity levels. In contrast, floxed control and *Piezo1* cKO groups remained below baseline levels, suggesting that *Piezo1* cKO alone does not restore activity changes after DMM surgery. These findings highlight the unique recovery of *Piezo2* cKO and *Piezo1/2* cKO mice, which achieved near-baseline running distances by 12 weeks.

Female mice were evaluated at the same timepoints (n = 6–15/genotype) (**Figure 5J, Supplemental Figure 6E-H**). At baseline (0 weeks), a one-way ANOVA confirmed no significant differences among groups, indicating similar activity levels before surgery (**Supplemental Figure 6E**). At 4 weeks post-DMM, a one-way ANOVA showed no significant differences in the distance run between the groups (P = 0.163) (**Supplemental Figure 6F**). At this time point, the floxed controls, *Piezo2* KO, and *Piezo1/2* cKO groups ran distances near or slightly below the baseline control values, while *Piezo1* cKO mice maintained distances above the baseline. At 8 weeks post-DMM, a one-way ANOVA detected no significant differences among groups (P = 0.298) (**Supplemental Figure 6G**). All genotypes exceeded or matched their baseline levels, showing little change in activity compared to pre-surgery distances. Similarly, at 12 weeks post-DMM, the one-way ANOVA showed no significant differences in distance run (P = 0.071) (**Figure 5J, Supplemental Figure 6H**). However, only the *Piezo1/2* cKO group exceeded the baseline control values, suggesting partial recovery of running activity to pre-surgery levels. Overall, voluntary running distances at the study endpoint remained comparable across genotypes for female mice.

### Structural Joint Changes are Correlated with Pain Behaviors in Male Mice Post-DMM Surgery

We investigated the relationships between pain behaviors and structural joint changes, grouping data by distinct behavioral measures across genotypes in male mice (**Table 1**). The Grimace Score, which reflects spontaneous pain,^28^ revealed significant correlations with multiple structural changes. In *Piezo2* cKO mice, a significant positive correlation linked Grimace Score to the Modified Mankin Score (P = 0.006, R^²^ = 0.62), showing that increased cartilage damage correlates with heightened pain perception. Similarly, *Piezo2* cKO mice demonstrated a significant positive correlation between osteophyte formation and Grimace Score (P = 0.049, R^²^ = 0.45), highlighting the association between joint changes and pain perception. In the *Piezo1* cKO and *Piezo1/2* cKO groups, significant positive correlations connected synovitis with Grimace Score (P = 0.032, R^²^ = 0.72 for *Piezo1* cKO; P = 0.059, R^²^ = 0.31 for *Piezo1/2* cKO), indicating that increasing inflammation is linked to heightened pain behaviors. The Electronic Von Frey (EVF), which measures mechanical sensitivity, showed notable correlations with the Modified Mankin Score in *Piezo1* cKO mice (P = 0.002, R^²^ = 0.93, negative trend). This finding suggests that greater cartilage damage leads to heightened sensitivity, indicating a worsening pain response as structural integrity declines. The SMALGO test showed a significant negative correlation with synovitis in *Piezo2* cKO mice (P = 0.037, R² = 0.54). This inverse relationship suggests that increased synovial inflammation may reduce mechanical hypersensitivity, implying a compensatory or altered pain response under inflammatory conditions. Interestingly, *Piezo1* cKO mice exhibited a significant positive correlation between Modified Mankin Score and Distance Run (P = 0.017, R² = 0.75). Despite increased joint damage, these mice maintained higher activity levels, suggesting compensatory behavior or reduced sensitivity to joint damage compared to other genotypes.

**Table 1.** Male Structural Changes Correlated with Pain and Behavior Assessments. This table summarizes the correlation analyses between structural changes (Modified Mankin Score, synovitis, and osteophyte scores) and various pain and behavior assessments (Grimace, SMALGO, EVF, SWB, and Distance) in male mice across floxed, *Piezo1* cKO, *Piezo2* cKO, and *Piezo1/2* cKO genotypes (N = 6-10/genotype). Each cell displays the P value and corresponding R² value for the correlation. Bold text denotes cells where the P value is less than 0.1, indicating a trend or significant correlation.

|  | Grimace |  | SMALGO |  | EVF |  | SWB |  | Distance |  |
| --- | --- | --- | --- | --- | --- | --- | --- | --- | --- | --- |
| Mankin | P | $R^2$ | P | $R^2$ | P | $R^2$ | P | $R^2$ | P | $R^2$ |
| Floxed | 0.556 | 0.03 | 0.321 | 0.08 | 0.986 | < 0.01 | 0.770 | 0.01 | 0.247 | 0.10 |
| *Piezo1* cKO | 0.169 | 0.41 | 0.435 | 0.15 | **0.001** | 0.93 | 0.195 | 0.37 | **0.016** | 0.74 |
| *Piezo2* cKO | **0.007** | 0.62 | 0.460 | 0.08 | 0.264 | 0.17 | 0.147 | 0.36 | 0.138 | 0.24 |
| *Piezo1/2*<br>cKO | 0.209 | 0.13 | 0.922 | < 0.01 | 0.681 | 0.02 | 0.809 | 0.01 | 0.334 | 0.12 |
| Synovitis | P | $R^2$ | P | $R^2$ | P | $R^2$ | P | $R^2$ | P | $R^2$ |
| Floxed | 0.155 | 0.15 | 0.061 | 0.26 | 0.606 | 0.02 | 0.784 | 0.78 | 0.273 | 0.03 |
| *Piezo1* cKO | **0.032** | 0.72 | 0.995 | < 0.01 | **0.003** | 0.91 | 0.105 | 0.52 | **0.072** | 0.51 |
| *Piezo2* cKO | 0.313 | 0.14 | **0.037** | 0.54 | 0.390 | 0.12 | 0.704 | 0.03 | 0.866 | 0.03 |
| *Piezo1/2* cKO | **0.059** | 0.31 | 0.123 | 0.24 | 0.289 | 0.28 | 0.867 | < 0.01 | 0.857 | 0.04 |
| **Osteophytes** | **P** | **R<sup>2</sup>** | **P** | **R<sup>2</sup>** | **P** | **R<sup>2</sup>** | **P** | **R<sup>2</sup>** | **P** | **R<sup>2</sup>** |
| Floxed | 0.661 | 0.02 | 0.286 | 0.10 | 0.688 | 0.01 | 0.715 | 0.02 | 0.466 | 0.03 |
| *Piezo1* cKO | **0.073** | 0.59 | 0.416 | 0.17 | 0.217 | 0.34 | 0.328 | 0.23 | 0.977 | < 0.01 |
| *Piezo2* cKO | **0.049** | 0.45 | 0.131 | 0.13 | 0.125 | 0.27 | 0.907 | < 0.01 | >0.999 | < 0.01 |
| *Piezo1/2* cKO | 0.820 | 0.82 | 0.114 | 0.28 | 0.926 | < 0.01 | 0.690 | 0.03 | 0.793 | 0.01 |

*Piezo2* cKO mice demonstrated the most consistent associations between structural joint changes – such as cartilage damage and osteophytes – and pain behavior measures, particularly Grimace Score and EVF. This indicates that cartilage degradation and osteophyte formation strongly influence pain perception in *Piezo2* cKO mice. In contrast, the *Piezo1* cKO genotype exhibited strong correlations involving synovitis, suggesting that inflammation plays a central role in pain perception in this group. Overall, *Piezo1/2* cKO and floxed control genotypes had fewer significant correlations, suggesting resilience to structural damage or compensatory mechanisms that reduce the impact of joint changes on pain behavior. These findings offer insights for developing osteoarthritis treatments by targeting specific structural features like cartilage damage, synovitis, and osteophytes to alleviate pain. Future research should prioritize interventions that prevent these structural changes, aiming to manage osteoarthritis-related pain more effectively.

### Female Mice Exhibit Genotype-Specific Links Between Joint Damage and Pain Behaviors

We analyzed correlations between structural joint changes and pain/behavior assessments in female mice, revealing distinct but generally less consistent patterns compared to male mice (Table 2). The *Piezo2* cKO genotype exhibited a significant negative correlation between the Modified Mankin Score and static weight-bearing (P = 0.022, R^²^ = 0.76). This indicates that increasing cartilage damage reduces weight-bearing capacity, reflecting a worsening functional outcome in response to joint degradation. Additionally, in *Piezo1/2* cKO mice, showed a trend toward a negative correlation between synovitis and SWB (P = 0.087, R^²^ = 0.36), suggesting that increased synovial inflammation may impair weight-bearing on the surgical limb. In the *Piezo1* cKO group, a trend toward significance linked osteophyte formation and Grimace Score (P = 0.078, R^²^ = 0.49). This trend implies that greater osteophyte formation correlates with heightened pain behaviors, highlighting a potential association between structural changes and pain perception. A significant positive correlation emerged between osteophyte formation and EVF threshold in both *Piezo1* cKO and *Piezo1/2* cKO mice (P = 0.063 for *Piezo1* cKO; P = 0.03 for *Piezo1/2* cKO; R^²^ = 0.53 and 0.35, respectively). This positive relationship suggests a complex or contradictory interaction between osteophyte formation and sensitivity to external mechanical stimuli. Overall, the *Piezo1* cKO and *Piezo2* cKO genotypes demonstrated significant correlations or trends linking structural damage (such as cartilage damage and osteophytes) to behavioral outcomes, indicating a higher sensitivity to joint damage compared to floxed control and *Piezo1/2* cKO genotypes. The floxed control and *Piezo1/2* cKO groups showed fewer significant correlations, suggesting greater resilience to joint changes or alternative compensatory mechanisms in female mice. These findings highlight potential sex-specific differences in joint degeneration and pain perception, which future research on osteoarthritis-related pain interventions should consider.

**Table 2.**
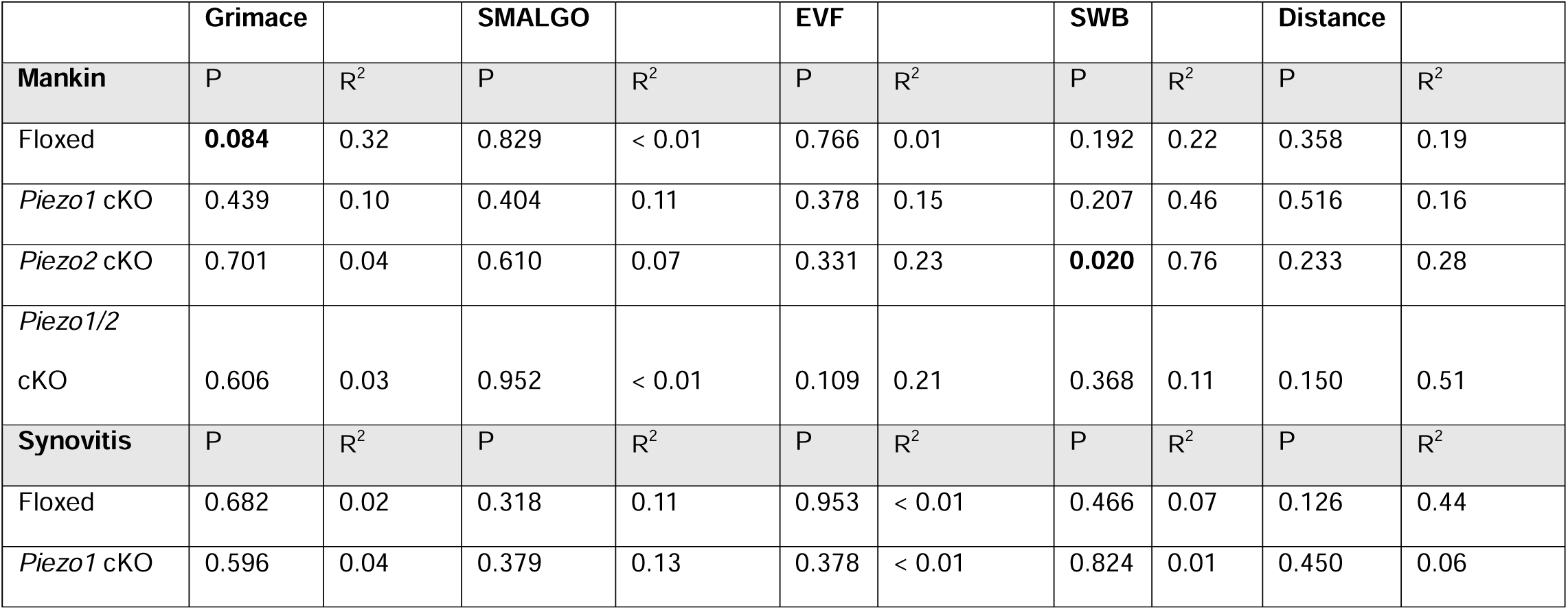
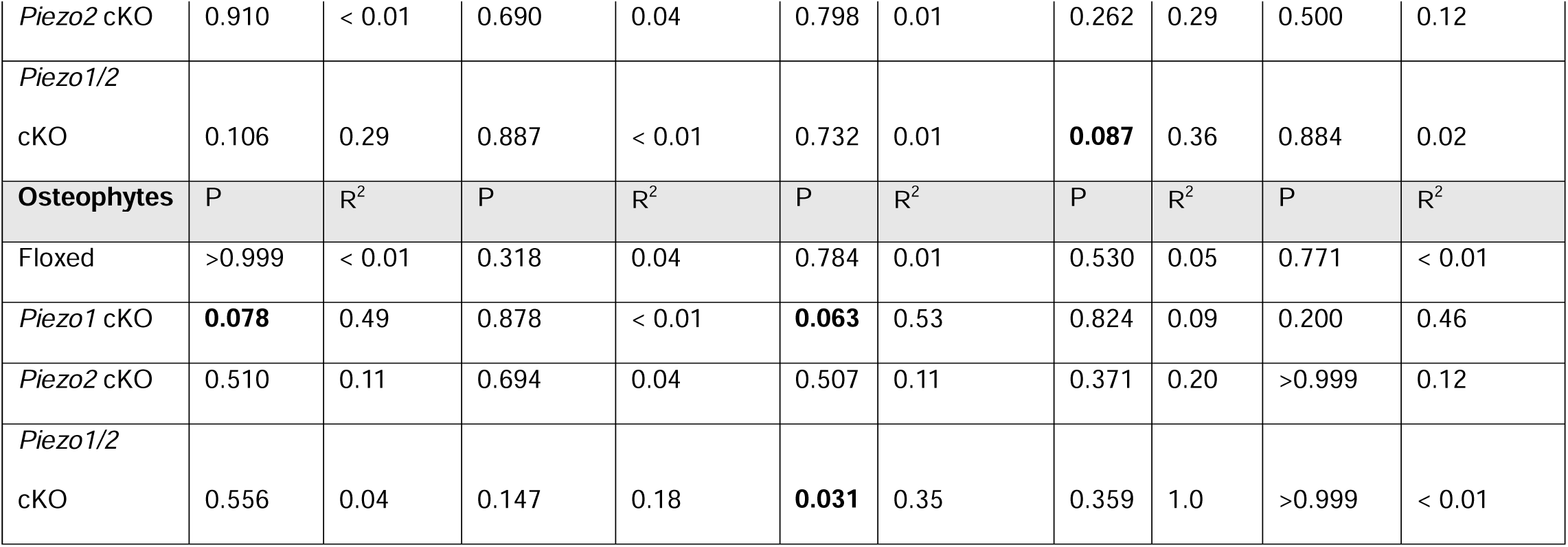
Female Structural Changes Correlated with Pain and Behavior Assessments. This table summarizes the correlation analyses between structural changes (Modified Mankin Score, synovitis, and osteophyte scores) and various pain and behavior assessments (Grimace, SMALGO, EVF, SWB, and Distance) in female mice across floxed, *Piezo1* cKO, *Piezo2* cKO, and *Piezo1/2* cKO genotypes (N = 6-10/genotype). Each cell displays the P value and corresponding R² value for the correlation. Bold text denotes cells where the P value is less than 0.1, indicating a trend or significant correlation. These analyses reveal genotype-specific relationships, providing insight into the interplay between structural joint changes and pain-related behaviors in female mice.

|  | Grimace |  | SMALGO |  | EVF |  | SWB |  | Distance |  |
| --- | --- | --- | --- | --- | --- | --- | --- | --- | --- | --- |
| **Mankin** | P | $R^2$ | P | $R^2$ | P | $R^2$ | P | $R^2$ | P | $R^2$ |
| Floxed | **0.084** | 0.32 | 0.829 | < 0.01 | 0.766 | 0.01 | 0.192 | 0.22 | 0.358 | 0.19 |
| *Piezo1* cKO | 0.439 | 0.10 | 0.404 | 0.11 | 0.378 | 0.15 | 0.207 | 0.46 | 0.516 | 0.16 |
| *Piezo2* cKO | 0.701 | 0.04 | 0.610 | 0.07 | 0.331 | 0.23 | **0.020** | 0.76 | 0.233 | 0.28 |
| *Piezo1/2*<br>cKO | 0.606 | 0.03 | 0.952 | < 0.01 | 0.109 | 0.21 | 0.368 | 0.11 | 0.150 | 0.51 |
| **Synovitis** | P | $R^2$ | P | $R^2$ | P | $R^2$ | P | $R^2$ | P | $R^2$ |
| Floxed | 0.682 | 0.02 | 0.318 | 0.11 | 0.953 | < 0.01 | 0.466 | 0.07 | 0.126 | 0.44 |
| *Piezo1* cKO | 0.596 | 0.04 | 0.379 | 0.13 | 0.378 | < 0.01 | 0.824 | 0.01 | 0.450 | 0.06 |
| *Piezo2* cKO | 0.910 | < 0.01 | 0.690 | 0.04 | 0.798 | 0.01 | 0.262 | 0.29 | 0.500 | 0.12 |
| *Piezo1/2*<br>cKO | 0.106 | 0.29 | 0.887 | < 0.01 | 0.732 | 0.01 | **0.087** | 0.36 | 0.884 | 0.02 |
| **Osteophytes** | **P** | **R<sup>2</sup>** | **P** | **R<sup>2</sup>** | **P** | **R<sup>2</sup>** | **P** | **R<sup>2</sup>** | **P** | **R<sup>2</sup>** |
| Floxed | >0.999 | < 0.01 | 0.318 | 0.04 | 0.784 | 0.01 | 0.530 | 0.05 | 0.771 | < 0.01 |
| *Piezo1* cKO | **0.078** | 0.49 | 0.878 | < 0.01 | **0.063** | 0.53 | 0.824 | 0.09 | 0.200 | 0.46 |
| *Piezo2* cKO | 0.510 | 0.11 | 0.694 | 0.04 | 0.507 | 0.11 | 0.371 | 0.20 | >0.999 | 0.12 |
| *Piezo1/2*<br>cKO | 0.556 | 0.04 | 0.147 | 0.18 | **0.031** | 0.35 | 0.359 | 1.0 | >0.999 | < 0.01 |

**Table 3.**
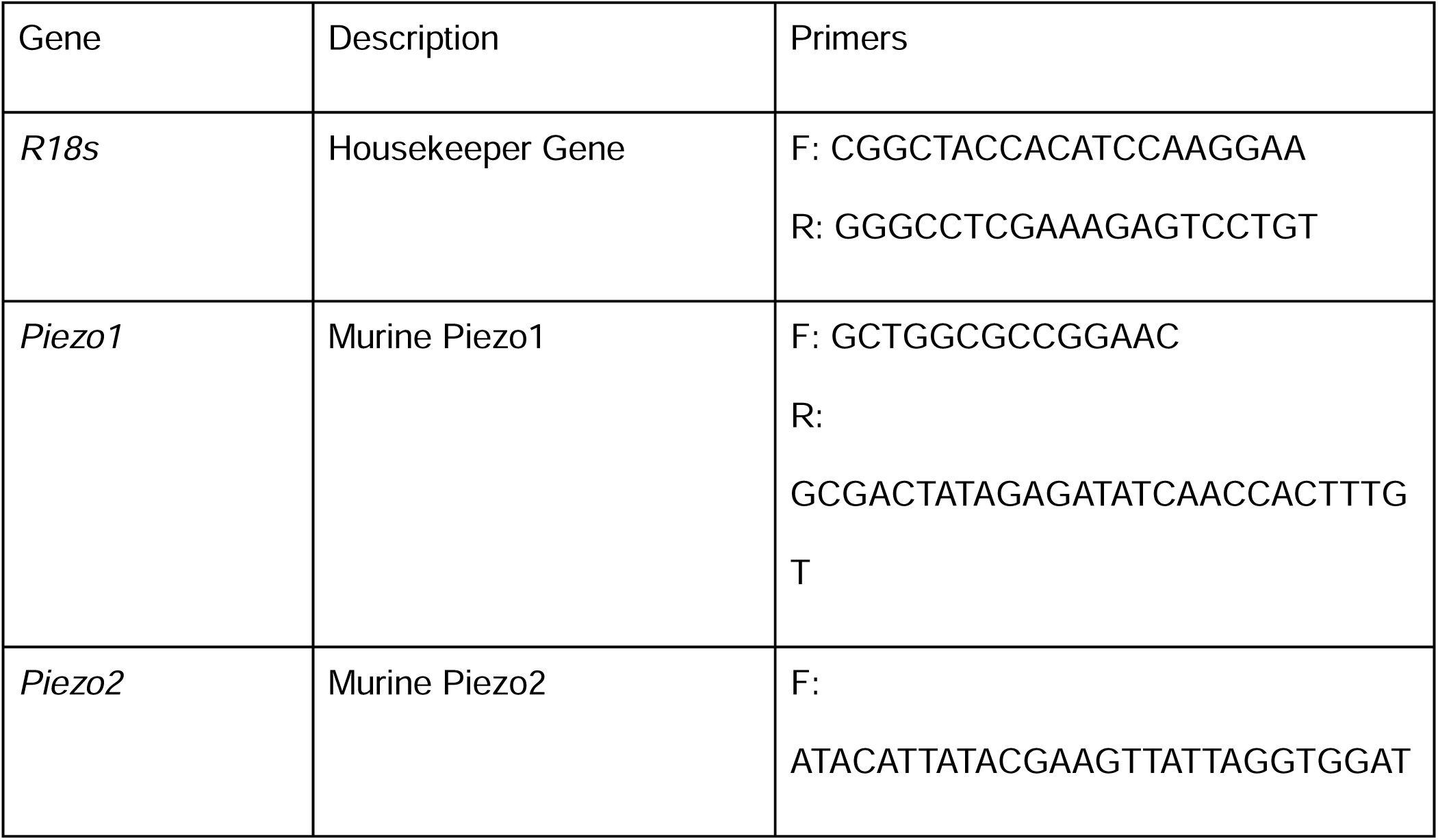
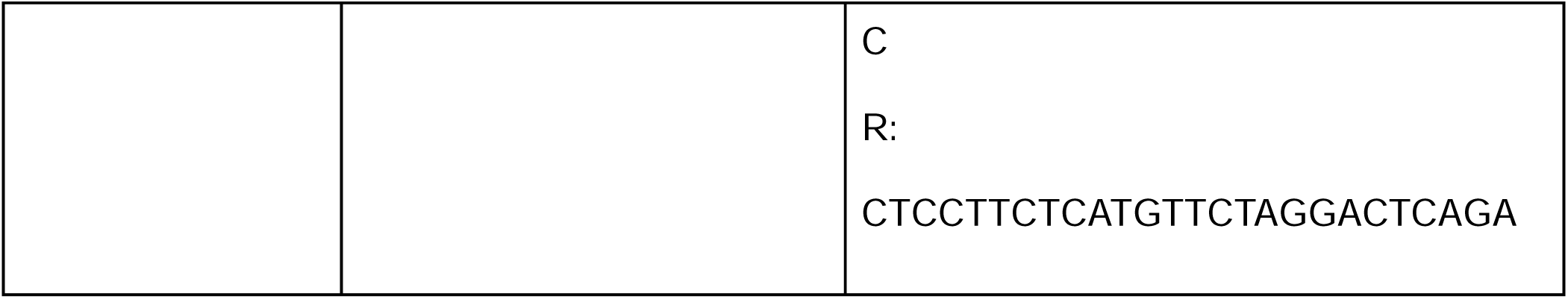
RT-PCR Primer sequence for Piezo Ion Channels.

## Discussion

In this study, we interrogated the independent and combinatory roles of the mechanosensitive ion channels Piezo1 and Piezo2 in post-traumatic osteoarthritis (OA) in male and female mice using an inducible, chondrocyte-specific, aggrecan-cre driver to knockout *Piezo1*, *Piezo2*, and both channels simultaneously in mice. Unlike prior studies that have focused on singular roles or shorter timelines for disease development, our work stands apart by providing a comprehensive analysis of these channels’ distinct and synergistic contributions to OA progression, pain, and inflammation over an extended disease timeline. Our findings support the hypothesis that targeting the Piezo ion channels can attenuate OA progression and pain, providing insights into their contributions to joint pathology and potential therapeutic value. *Piezo1* and *Piezo2* demonstrated distinct and complementary roles in mediating cartilage integrity, inflammation, and pain behaviors. While *Piezo1* deletion delayed the onset of pain progression, it ultimately resulted in severe cartilage degradation and heightened synovial inflammation, suggesting a dual role in both protective and pathogenic mechanotransduction. *Piezo2* knockout, on the other hand, reduced pain behaviors but was not protective against cartilage damage in male and female mice. Most notably, the combined deletion of *Piezo1* and *Piezo2* provided protection against cartilage degeneration, synovitis, spontaneous pain, hyperalgesia, changes in hindlimb loading, and spontaneous activity, highlighting the synergistic functions of these channels and the power of targeting their overlapping mechanosensitive functions. Together, these results underscore the importance of Piezo ion channels in OA pathogenesis and support their individual and combinatory potential as therapeutic targets for addressing both structural and symptomatic aspects of the disease.

Our findings highlight the protective effect of combined deletion of *Piezo1* and Piezo2 in osteoarthritis (OA) progression, as evidenced by significantly lower Modified Mankin scores in the *Piezo1/2* cKO group compared to both the control and single-knockout groups. This protection was observed in both male and female mice, with an additional sex-specific observation in female *Piezo1/2* cKO mice, where the DMM limb did not develop significantly higher Modified Mankin scores than the contralateral limb. While the deletion of either *Piezo1* or *Piezo2* alone may lead to partial impairment in mechanical signaling, the presence of the other may be sufficient to sustain disease-driving mechanotransductive pathways.^14^ These results suggest that the Piezo1 and Piezo2 ion channels have overlapping functions and that eliminating both is necessary to mitigate the pathogenic mechanosensitive response in chondrocytes, aligning with prior findings that both channels contribute to cartilage health under physiological loading.^19^

While previous studies demonstrated that *Piezo1* knockout alone is protective against OA onset,^18,24-26^ our findings challenge the current paradigm by showing that *Piezo1* deletion, while possibly delaying pain progression, does not prevent severe cartilage degradation or inflammation by 12 weeks post-DMM surgery. Complementing previous studies that employed the ACLT model^24,25^ or an 8-week assessment period,^26^ we evaluated OA progression at 12 weeks post-DMM surgery, a longer timeline that may reveal late-stage disease phenotypes. Additionally, our use of an aggrecan-cre driver, with induction at 12 weeks of age, to achieve chondrocyte-specific knockouts may differ from the *Col2a1*-cre^25^ or *Gdf5-*cre^23^ drivers employed by other groups. For example, *Col2a1* expression decreases significantly after joint development in mice, and both *Col2a1* and *Gdf5* cre drivers may be expressed in other joint tissues.^29,30^ Together, these differences emphasize the importance of study design in interpreting the role of Piezo1 ion channels in OA. While *Piezo1* deletion may delay the onset of cartilage degeneration, our findings suggest that it is insufficient to prevent long-term structural damage and pain phenotypes.

The *Piezo2* cKO genotype demonstrated the most severe cartilage damage in female mice, although not significant, as reflected by increased Modified Mankin scores. This observation is consistent with a prior study using the ACLT model, which also reported a lack of chondroprotection in *Piezo2* cKO female mice.^25^ The increased cartilage degradation in *Piezo2* cKO mice may be partially attributed to compensatory upregulation and activation of *Piezo1*, as evidenced by our qPCR data. These findings underscore the distinct and sex-specific roles of *Piezo2* in cartilage maintenance and suggest that the absence of *Piezo2,* coupled with active *Piezo1*, may worsen OA pathology.

Interestingly, osteophyte formation was most pronounced in *Piezo1* cKO male mice in our study. Although our findings were not significant, they provide an interesting contrast with findings from a previous study that reported a significant reduction in osteophyte size with *Piezo1* knockout using *Col2a1-*Cre mice.^25^ The differences between these findings may stem from variations in experimental design and evaluation methods. Brylka et al. utilized the ACLT model, assessed osteophytes at an earlier 8-week timepoint, and employed micro-computed tomography (µCT) to evaluate osteophyte formation.^25^ Information on their cre induction is not accessible so we are unable to compare to our methods. In contrast, our study used the DMM model, evaluated osteophytes at 12 weeks post-surgery, and relied on histological assessments, which may capture different aspects of osteophyte development. Our study also observed reduced osteophyte numbers in *Piezo2* cKO mice, further underscoring the distinct roles of *Piezo1* and *Piezo2* in joint remodeling. Notably, *Piezo1/*2 cKO mice showed no significant increase in osteophyte formation when comparing the contralateral versus the DMM limb. This suggests that the deletion of *Piezo1* and *Piezo2* may mitigate osteophyte formation, possibly through the reduction of mechanosensitive signaling pathways involved in pathological bone remodeling. It is worth noting that female mice osteophyte numbers were consistent independent of genotype and surgery. These findings provide new insights into the roles of Piezo channels in regulating joint remodeling and highlight the importance of experimental context in interpreting osteophyte outcomes.

Our results revealed that synovitis scores were significantly elevated in the DMM limbs compared to contralateral limbs across all genotypes, except in the *Piezo1/2* cKO group. This suggests that the simultaneous deletion of *Piezo1* and *Piezo2* mitigates synovial inflammation, unlike the single knockouts where inflammation was prominent. Both *Piezo1* and *Piezo2* cKO groups displayed heightened synovitis scores, highlighting their individual contributions to the inflammatory response following joint injury. The elevated synovitis observed in *Piezo2* cKO mice corresponds with their high Modified Mankin scores, further implicating *Piezo1* in sensitizing chondrocytes to inflammatory signaling and promoting a feed-forward mechanism that exacerbates OA progression.^17^ This mechanism may explain the increased synovial inflammation in the *Piezo2* cKO group, where compensatory upregulation and activation of *Piezo1* likely amplify the inflammatory response. In contrast, the *Piezo1/2* cKO group exhibited no significant synovitis in their DMM limbs compared to contralateral limbs, suggesting that eliminating both channels disrupts the pathogenic signaling cascade and reduces synovial inflammation. Although cre recombination in synovial tissue is unlikely due to the absence of aggrecan expression in synovial fibroblasts or macrophages under normal conditions, our findings support a role for Piezo channels in modulating synovial inflammation, likely through cartilage-derived signaling pathways. The reduced synovitis scores in *Piezo1/2* cKO mice, compared to elevated inflammation in single knockouts, indicate a functional role for Piezo-mediated mechanotransduction in driving synovial responses following joint injury. Additionally, while our model does not permit direct interrogation of Piezo function in synovial cells, recent studies have demonstrated that *Piezo1* is expressed in the synovium and may regulate inflammatory signaling and cell proliferation in response to mechanical loading.^3,31^ Exploring Piezo channel function specifically in the synovium remains an important future direction for understanding their contribution to joint inflammation.

Our µCT analysis highlights the critical role of *Piezo2* in bone structure and remodeling, with significant changes observed in bone volume/total volume (BV/TV), trabecular thickness, trabecular number, bone mineral density (BMD), medial tibial plateau thickness, and meniscus ossification in both contralateral and DMM limbs in male and female mice. These findings suggest that *Piezo2* is essential for maintaining bone integrity, independent of joint injury or trauma. Interestingly, *Piezo1/2* cKO mice displayed reduced medial tibial plateau thickness in both sexes, and female *Piezo1/2* cKO mice also exhibited decreased BMD, further emphasizing the importance of *Piezo2* in bone remodeling processes.

The importance of Piezo channels in bone integrity is consistent with findings from previous studies. For example, Brylka et al. observed that *Piezo1*-Col2a1Cre mice exhibited a marked reduction in subchondral bone volume and trabecular bone, largely attributable to a decrease in trabecular number.^25^ However, these effects were not observed in *Piezo2*-Col2a1Cre mice, suggesting distinct roles for *Piezo1* and *Piezo2* in bone remodeling. Unlike the Brylka study, our analysis did not detect significant changes in bone structure in *Piezo1* cKO mice, potentially due to differences in genetic drivers, injury models, or assessment methods, which we describe in more detail above. While the role of *Piezo2* in bone biology remains largely unexplored, our findings are the first to identify *Piezo2* as a key regulator of bone remodeling. This work also expands on existing literature that has firmly established *Piezo1* as essential for bone growth and maintenance.^32,33^

The distinct reductions in BV/TV, BMD, trabecular structure, tibial plateau thickness, and meniscal ossification, independent of surgery, in *Piezo2* cKO mice may stem from impaired mechanotransduction in chondrocytes and other aggrecan-expressing cells, such as meniscus or bone progenitors, disrupting their ability to adapt to mechanical forces. This imbalance likely affects endochondral ossification, resulting in reduced bone structure and mineral density. Our data rule out changes in Piezo expression in the growth plate as a driving factor of the observed structural changes. These data underscore the importance of chondrocyte *Piezo2* in regulating the adaptive response of bone to mechanical loading and highlight its potential as a therapeutic target for addressing bone changes associated with OA progression. Furthermore, as bone remodeling may influenced by altered joint loading secondary to pain responses, the effects of *Piezo* on bone may occur through multiple potential pathways. The findings also suggest that dual inhibition of *Piezo*1 and *Piezo2* may further alter bone remodeling, necessitating careful consideration in the context of therapeutic development.

The assessment of pain and behavior outcomes, including grimace scores, pressure-pain hyperalgesia thresholds, and voluntary running activity, revealed distinct, genotype-dependent responses following DMM surgery. Both *Piezo2* cKO and *Piezo1/2* cKO mice exhibited reduced pain behaviors, suggesting that *Piezo2* plays a significant role in pain perception and sensitization, consistent with its established role in mechanosensation and pain signaling in nociceptors.^22^ Despite severe cartilage degradation, synovial inflammation, and bone structural changes, *Piezo2* cKO mice demonstrated lower pain scores and higher activity levels, while *Piezo1/*2 cKO mice showed the greatest reduction in pain, with grimace scores and hyperalgesia thresholds approaching baseline levels by 12 weeks post-DMM, indicating a potential protective effect against pain. In contrast, *Piezo1* cKO mice exhibited increased pain scores and reduced activity, particularly in female mice, suggesting that *Piezo1* may contribute to persistent pain after joint injury.

Our time course analysis provided additional insights into OA pain progression, showing that changes in grimace scores and pain emerged as early as 4 weeks post-surgery in female mice, while significant changes in male mice were noted only at 12 weeks. Similarly, pressure-pain hyperalgesia and tactile allodynia appeared at 4 weeks in female mice and 8 weeks in males. Load distribution changes were observed by 4 weeks in both sexes, but only *Piezo1/2* cKO males achieved complete resolution by 12 weeks. Changes in wheel running activity emerged at 12 weeks in both sexes. These findings highlight the importance of extended pain and activity measurements, supporting a minimum 12-week timeline to fully capture OA pain progression, consistent with prior studies.^22,34,35^ This study is among the first to comprehensively evaluate how simultaneous targeting of *Piezo1* and *Piezo2* impacts both structural and pain outcomes, providing a dual-target therapeutic rationale for OA management.

Correlations between structural changes and pain behaviors highlighted the therapeutic relevance of addressing specific OA-related features. In male mice, these correlations emphasized the importance of targeting cartilage damage, synovitis, and osteophyte formation to alleviate pain, particularly in *Piezo2* cKO mice. In *Piezo1* cKO mice, synovitis appeared to play a central role in pain outcomes, suggesting that anti-inflammatory therapies could be effective in mitigating synovitis-driven pain. Conversely, the fewer correlations observed in female mice point to potential sex-specific biological mechanisms underlying degeneration and pain, underscoring the necessity of tailored treatment strategies. Interestingly, the *Piezo1/2* cKO genotype exhibited fewer correlations overall, suggesting the presence of compensatory mechanisms that may reduce the impact of joint damage. These findings underscore the potential for interventions that simultaneously address cartilage integrity, inflammation, and bone remodeling, while considering patient-specific differences, to enhance OA pain management.

The observed sex-specific differences in cartilage damage and pain outcomes further highlight the distinct roles of Piezo channels in male and female joints, emphasizing the need for tailored therapeutic strategies. For instance, the heightened cartilage damage in female *Piezo2* cKO mice suggests that *Piezo2*-targeted therapies may require sex-specific optimization to achieve maximum efficacy.

Our findings underscore the potential therapeutic value of targeting both *Piezo1* and *Piezo2* in OA. While *Piezo1* inhibition alone did not prevent cartilage degradation or synovial inflammation, it may delay the onset and progression of pain and structural damage. Conversely, *Piezo2* inhibition reduced pain and preserved activity levels but did not impact structural damage, potentially due to compensatory *Piezo1* activation. The dual inhibition observed in *Piezo1/*2 cKO mice suggests that simultaneously targeting both channels could effectively mitigate cartilage damage and pain. These results support the development of Piezo channel inhibitors as promising disease-modifying OA drugs that address both structural and symptomatic aspects of the disease.

While our study provides critical insights into the roles of Piezo1 and Piezo2 ion channels in OA, several limitations warrant consideration. First, we focused on a single surgical model of OA; validating these findings across additional injury and degeneration models would strengthen their generalizability. Second, we did not include sham-operated controls, which limits our ability to distinguish the short-term effects of surgery from those of DMM-induced degeneration. While previous studies have shown that sham surgery does not lead to OA in this model^34,36^, including sham groups in future studies would help isolate the specific contributions of joint destabilization to the observed outcomes. Additionally, the use of genetic ablation models limits the translational relevance to pharmacological interventions. Future studies could address this by using pharmacological inhibitors or siRNA approaches to assess whether similar protective effects are observed. The precise mechanisms by which *Piezo1* and *Piezo2* influence synovitis, osteophyte formation, and pain also remain unclear. Investigating downstream signaling pathways in chondrocytes and synoviocytes will be essential for understanding their roles in OA pathogenesis. For example, exploring the link between Piezo channel activation and chondrocyte-mediated senescence could shed light on the mechanisms underlying cartilage protection in double knockout mice.

An important point to consider in the interpretation of our study is that aggrecan is expressed in other tissues besides cartilage, such as perineuronal nets and dorsal root ganglia. We observed changes in RNA expression in the lung and brain in the *Piezo2* and *Piezo1* cKO mice, highlighting the importance of considering these off-target effects moving forward. As a result, the pain and behavior phenotypes observed, particularly in the *Piezo2* cKO mice, may be influenced by off-target effects of aggrecan knockout in non-cartilage tissues. This broader expression could contribute to the altered pain sensitivity or behavior we observed, and future studies should explore other tissue-specific knockouts or different knockout mechanisms to better isolate the role of Piezo channels in cartilage.

In summary, this work provides a multi-factorial and sex-specific analysis of *Piezo1* and *Piezo2’*s roles in OA using innovate study designs to uncover findings that challenge existing paradigms in the field. Our comprehensive integration of structural, inflammatory, and pain data offers a unique insight of these channels’ contributions, paving the way for targeted, disease-modifying therapeutics. Our findings suggest that *Piezo1* and *Piezo2* have overlapping and compensatory functions in maintaining joint health and understanding their interplay and integration with other mechanosensitive pathways, such as TRPV4, will be crucial for developing targeted therapies that modulate mechanotransduction in OA without compromising normal joint function.

## Methods

### Generation of Conditional Knockout Mice

All procedures were conducted in accordance with protocols approved by the Washington University in St. Louis Institutional Animal Care and Use Committee. To generate conditional knockout mice, (Agc)1^tm(IRES-CreERT2)^ mice (B6.Cg-Acan^tm1(cre/ERT2)Crm^/J, Jackson strain #019148) were crossed with Piezo1^fl/fl^ mice (B6.Cg-Piezo1^tm2.1Apat^/J, Jackson strain #029213) and Piezo2^fl/fl^ (B6(SJL)-Piezo2^tm2.2Apat^/J, Jackson strain #027720) mice to generate (Agc)1-CRE^ERT2^;Piezo1^fl/fl^ (*Piezo*1 cKO), (Agc)1-CRE^ERT2^;Piezo2^fl/fl^ (*Piezo*2 cKO), (Agc)1-CRE^ERT2^;Piezo1^fl/fl^ Piezo2^fl/fl^ (*Piezo*1/2 cKO), and Piezo1^fl/fl^ Piezo2^fl/fl^ (floxed) mice. Cre-negative littermates (floxed), served as controls. Genotyping was performed by Transnetyx (Cordova, TN). Following weaning, all animals were fed a 10% fat chow diet (PicoLab Rodent Diet 5053, LabDiet). Both male and female mice were included (n=10-15/genotype/sex). Tamoxifen (Sigma, St. Louis, MO, T5648) was administered intraperitoneally (100 mg/kg body weight in corn oil) for 5 consecutive days to induce Cre-mediated recombination, starting at 12 weeks of age. Both floxed control and transgenic mice received Tamoxifen. Mice were housed randomly with littermates in groups of 3–4 per cage. Functional Ca^2+^ imaging on isolated femoral condyles was performed two weeks post-induction, and further analyses were conducted 16 weeks post-induction.

### Quantitative Polymerase Chain Reaction

At 28 weeks of age, floxed, *Piezo1* cKO, *Piezo2* cKO, and *Piezo1/2* cKO mice were euthanized, and the right hip cap was immediately harvested (n=6-10/genotype). The tissue was placed in RL buffer and processed for RNA isolation according to the manufacturer’s protocol (48300; Norgen Biotek, Thorold, ON, Canada). Reverse transcription was performed using the SuperScript VILO cDNA synthesis kit (11755500; Life Technologies, Carlsbad, CA, USA) following the manufacturer’s instructions. Quantitative polymerase chain reaction (qPCR) was conducted using FASTSybr (4385617; Applied Biosystems, Waltham, MA, USA) according to the manufacturer’s recommendations. The cycling conditions were: an initial denaturation at 95°C for 10 minutes, followed by 40 cycles of 95°C for 15 seconds, and 60°C for 60 seconds for annealing and extension. Gene expression fold changes were calculated relative to the floxed control group, using 18S ribosomal RNA as a reference gene. Data are reported as fold changes and calculated using the 2^−ΔΔCt^ method. Statistical analysis was performed using a one-way ANOVA with Tukey’s multiple comparisons test. Primer pairs for *Piezo1* and *Piezo2* were synthesized by Integrated DNA Technologies, Inc.

### Immunohistochemistry

Murine cartilage sections were prepared for immunohistochemistry (IHC) using the following protocol. Slides were baked at 60°C for 1 hour and allowed to cool before staining. Sections were rehydrated by washing in xylene three times (5 minutes each), followed by 100% ethanol (2 minutes), 50% ethanol (2 minutes), and two washes in distilled water (2 minutes each). Boundaries around each section were drawn using a hydrophobic pen, and HistoReveal (Abcam, Cambridge, UK, ab103720) was applied for 5 minutes. Slides were then washed twice in 1X PBS (5 minutes each). A peroxidase blocking solution (3% hydrogen peroxide in methanol) was prepared and applied to the slides for 30 minutes. Following another two washes in 1X PBS, 2.5% goat serum (Vector Labs, #S-1012) was distributed to the sections and incubated for 30 minutes to 1 hour. The PIEZO1 (Novus Biologicals, Centennial, CO, 78537) and PIEZO2 (Novus Biologicals, 78624) primary polyclonal antibodies were then added, and slides were incubated overnight in a cold room. After aspirating the primary antibody, sections were washed three times with 1X PBS and 0.1% Tween 20 (5 minutes each). The secondary antibody (ImmPRESS® HRP Goat Anti-Rabbit IgG Polymer Detection Kit, MP-7451) was diluted as per the manufacturer’s instructions and incubated on the sections for 45 minutes. Following this, slides were washed three times in PBS and 0.1% Tween, then briefly rinsed with distilled water. DAB Substrate Kit, Peroxidase (HRP), with Nickel, (3,3’-diaminobenzidine) (Vector, SK-4100) was prepared according to kit instructions and applied to sections for 7 minutes, followed by a rinse with water. Hematoxylin was used for counterstaining, applied for 3 minutes and rinsed in water. Finally, slides were dehydrated through sequential washes in water, 50% ethanol, and 100% ethanol, followed by a dip in xylene. Slides were then mounted using Permount and glass cover slips and allowed to dry overnight in the hood. All sections were imaged using an Olympus VS120 high-resolution slide scanner.

### Calcium Imaging

Floxed, *Piezo1* cKO, *Piezo2* cKO, and *Piezo1/2* cKO mice were euthanized via CO₂ exposure followed by cervical dislocation. Hindlimbs were carefully dislocated, and femora were isolated by removing the surrounding muscle, ligament, and tendon tissues (n= 8-10). A custom imaging rig was designed to stabilize the femoral condyles at a 10° angle at the bottom of a 35 mL well, exposing the lower portion of the condyles for imaging (**Supplemental Figure 7**). The rig was secured to the well using adhesive (E6000, Eclectic), and the cut portion of the femur was adhered to the rig with the same adhesive. The samples were stained in staining solution 40 minutes prior to imaging. The staining solution consisted of 1.5 mL HBSS with 2.5% HEPES, 15 μL Sulfinpyrazone, 10 μL Fura Red prepared in 10 μL Pluronic Acid, and 10 μL Fluo4 prepared in 10 μL Pluronic Acid. After aspiration of the staining solution, 1 mL of imaging buffer (HBSS with 2.5% HEPES) was added to the well. Calcium imaging was performed using a Zeiss Confocal Microscope equipped with a perfusion system. The femoral condyles were imaged from below the perfusion chamber using a 10x objective and a 488 nm laser to capture Fluo4 and Fura Red signals. The imaging setup included a temperature-controlled environment set to 37°C. A syringe was used to manually introduce 20 µM Yoda1, a Piezo1 channel activator (5586, Tocris), perfusion solution into the chamber after 160 seconds of baseline imaging. Transmitted light imaging provided visualization of the condyles to track calcium signaling indicative of Piezo channel activation.

### Calcium Signaling Data Analysis

Image analysis was conducted using a custom ImageJ macro to segment individual chondrocytes from the Fluo4 channel. A maximum Z-projection of Fluo4 images was used to generate a mask, which was then applied to the original images to quantify the mean brightness of each chondrocyte. Data from the Fura Red channel were processed similarly. Quantitative analysis was performed using an R script. Fluorescence intensity was normalized to the baseline average fluorescence for each cell. Cells were classified as responders if the normalized brightness after Yoda1 perfusion exceeded the baseline brightness by more than 0.5 times the standard deviation. The percentage of responding cells was calculated for each sample by dividing the number of responsive cells by the total number of cells analyzed. Samples in which the joint shifted during imaging were excluded from analysis due to segmentation inaccuracies. Statistical analysis was performed using one-way ANOVA to assess differences between groups.

### Destabilization of the Medial Meniscus Surgery

Following the 3-week tamoxifen washout period, mice underwent Destabilization of the Medial Meniscus (DMM) surgery on the left limb at 16 weeks of age, with the contralateral limb serving as a control.^34,37-40^ Mice were anesthetized using isoflurane and maintained on heating pads for thermoregulation. Perioperative analgesia was provided with a single dose of buprenorphine sustained release formula (1 mg/kg). Hair was removed around the surgical site, and the area was prepared aseptically with alternating iodine and 70% alcohol scrubs, followed by sterile draping. A 3 mm longitudinal incision was made from the distal patella to the proximal tibial plateau. The joint capsule was incised along the medial patellar tendon, and blunt dissection of the fat pad was performed to expose the medial meniscotibial ligament. The ligament was transected, resulting in destabilization of the medial meniscus and increased mechanical stress on the joint. The joint capsule was sutured with 8-0 Vicryl (Johnson & Johnson, J401G), and the skin was closed with tissue adhesive. The animals were monitored for three days to ensure wound healing. Animals were taken out to 28 weeks of age, 12 weeks post-DMM surgery, as the end point.

### MicroCT Analysis

Following sacrifice, knee joints were fixed in 4% paraformaldehyde (PFA) for 48 hours. Fixed joints were scanned using the Bruker Skyscan 1176 and 1276 microCT systems. Samples were loaded onto a 20 mm bed and scanned with the following parameters: 0.5 mm aluminum filter, 9 µm resolution, and a binning of 2048 x 2048 pixels. Flatfield correction was performed prior to scanning to ensure image quality. Scout scans were used to determine the start and end positions for each sample scan, and scans were added to a batch for automatic execution. Scans were acquired, and data were subsequently processed for quantitative analysis. Scans were reconstructed using NRecon software (Bruker). Scan files were loaded, and dynamic range was set to 0–0.1. Circular regions of interest (ROIs) were defined around the samples. Reconstruction settings included a beam hardening correction of 25 and a ring artifact correction of 4. Misalignment was adjusted as needed to minimize artifacts. The medial and lateral tibial plateau regions were defined in CTAn, using the region from the end of the tibia up to the growth plate. The analysis included the following key steps: thresholding to segment cortical bone, despeckling to remove noise, shrink-wrapping to define the region of interest, and various morphological operations to refine the segmentation and remove gaps. Bitwise operations were used to select the bone cavity and exclude background. Finally, a histogram analysis was performed to determine bone mineral density (BMD) within the region of interest, and a 2D analysis was used for numerical measurements. Additionally, a custom processing task list was created to isolate and measure the medial, lateral, and total ossified menisci of the joints. The outcomes reported in this study included bone volume/total volume (BV/TV), bone surface/volume (BS/BV), trabecular thickness, trabecular number, trabecular separation, and bone mineral density. Lastly, the thickness of the entire medial and lateral tibial plateau region was measured and reported in the study.

### Histological Assessments of Joint Damage

Following microCT scanning, knee joints were decalcified for 48 hours using Cal-Ex II Decalcifier Solution (Fisher Scientific, CS511), dehydrated, and embedded in paraffin using an automated tissue processor (Leica Microsystems, ASP300S). Coronal sections (5 μm) were cut using a microtome and baked at 60°C for one hour to ensure adherence to the slides. Sections were stained with Safranin-O (Sigma-Aldrich, HT904-8FOZ) and Fast Green (Electron Microscopy Sciences, #15500) to quantify cartilage degeneration according to the OARSI histopathology standards, using the Modified Mankin Criteria.^41^ Additional sections were stained with Hematoxylin (Fisher Scientific, NC9064721) and Eosin B (Sigma-Aldrich, #2853) to evaluate synovitis, scored based on the Krenn criteria.^27^ All sections were imaged using an Olympus VS120 high-resolution slide scanner. Joint damage (Modified Mankin score) (n=8-10/genotype), synovitis (n=9-20/genotype), and osteophyte (n=7-15/genotype) formation were scored by three blinded graders, and the average score from the three graders is reported in this study.

### Mouse Grimace Scale (MGS) Assessment

Pain-related behavior in male and female floxed control, *Piezo1* cKO, *Piezo2* cKO, and *Piezo1/2* cKO mice was assessed using the Mouse Grimace Scale (MGS), a reliable and non-invasive method to evaluate spontaneous pain (n=6-15/genotype/sex). The assessment followed the established guidelines as described by Langford et al.^28^ Mice were observed in their home cages without disturbance at the same time each day. Two blinded scorers independently assessed the mice, and scores were averaged for analysis. The MGS evaluates key facial features, including orbital tightening, nose bulge, cheek bulge, ear position, and whisker change, which are combined into a comprehensive score to quantify pain severity. Higher scores indicate more severe pain-related behaviors. Assessments were performed prior to the DMM surgery and at 4-, 8-, and 12-weeks post-surgery. Statistical analysis was performed using a 1-way ANOVA to assess differences between groups.

### Pressure-Pain Hyperalgesia Assay

Male and female floxed control, *Piezo1* cKO, *Piezo2* cKO, and *Piezo1/2* cKO mice were used for the study. Pressure-pain hyperalgesia was assessed in both limbs using the Small Animal Algometer (SMALGO) device (Bioseb) (n=6-15/genotype/sex). Pain measurements were taken prior to destabilization of the medial meniscus (DMM) surgery and at 4-, 8-, and 12-weeks post-surgery before sacrifice. Both the surgical and contralateral limbs were evaluated to assess pain sensitivity. To prevent tissue damage, a maximum threshold of 450 g was set during testing^42^. The threshold for mouse tolerance was recorded for each measurement. For each limb, three pressure measurements were taken by an individual blinded to the genotypes, and the results were averaged to provide a representative score for analysis. Statistical analysis was performed using a 2-way ANOVA to assess differences between limbs at each timepoint.^40^

### Electronic Von Frey (EVF)

Male and female floxed control, *Piezo1* cKO, *Piezo2* cKO, and *Piezo1/2* cKO mice were used in this study to measure tactile allodynia. Tactile sensitivity was assessed in both limbs using the Electronic Von Frey (EVF) device (Bioseb) (n=6-15/genotype/sex). Pain measurements were conducted prior to destabilization of the medial meniscus (DMM) surgery and at 4-, 8-, and 12-weeks post-surgery, with the 12-week time point corresponding to the sacrifice of the animals. Both the surgical and contralateral limbs were evaluated to determine sensitivity to tactile stimuli. The threshold for tactile response was recorded for each measurement. For each limb, three measurements were taken by an individual blinded to the genotypes, and the results were averaged to provide a representative score for analysis. Statistical analysis was performed using a 2-way ANOVA to assess differences between limbs at each time point.

### Static Weight Bearing

Static weight bearing was assessed in male and female floxed control, *Piezo1* cKO, *Piezo2* cKO, and *Piezo1/2* cKO mice, before surgery and 4-, 8-, and 12-weeks post DMM (n=6-15/genotype/sex). Mice were weighed prior to the static weight bearing assessment to calculate the percentage of force relative to body weight. For the assessment, mice were positioned in the static weight bearing holder (Bioseb, EB2-BIO-SWB-M) with their front paws placed on the slanted edge and their hind paws positioned flat on the bottom force plates. Once each animal was properly positioned, the weight placed on each hind limb was recorded. For each mouse, three recordings were taken, by an individual blinding to genotypes, and the results were averaged for subsequent analysis. In this study, the percent force of each limb relative to body weight was analyzed, and statistical comparisons were made between the experimental groups using a 2-way ANOVA.

### Spontaneous Activity Wheel Assessment

Voluntary activity was assessed using wheel running in male and female floxed control, *Piezo1* cKO, *Piezo2* cKO, and *Piezo1/2* cKO mice before surgery and at 4, 8, and 12 weeks post-DMM (n=6-15/genotype/sex). Mice were housed individually in cages with unlimited access to a running wheel (Bioseb, BIO-ACTIVW). To allow for acclimatization, mice were placed in the cages 1 hour before the assessment began. A minimum threshold of 0.3 m was set prior to starting the assessment, which began at the start of the dark cycle and continued for 18 hours. Distance run was recorded and analyzed in this study. Statistical comparisons between experimental groups at each timepoint were performed using a 2-way ANOVA.

### Correlation Analysis Between Structural and Pain Assessments

To explore the relationship between structural joint changes and pain/behavior outcomes, correlation analyses were conducted for both male and female mice (n = 6-10 per group for each assessment). Structural measurements included the Modified Mankin Score, synovitis, and osteophyte scores, which were correlated with pain and behavior assessments: Electronic Von Frey (EVF), SMALGO (mechanical hypersensitivity), grimace score, percentage of load on the DMM limb, and distance run. The analyses were performed using GraphPad Prism software. For each correlation, we calculated the P value to determine statistical significance. If the P value was below the significance threshold (P = 0.05), the R² value was reported to quantify the strength and direction of the relationship between structural and pain assessments. Data from both male and female mice were analyzed to investigate potential sex-specific differences in correlations between structural joint degeneration and pain-related behavior.

### Statistical Analysis

Graphing and analysis were performed using GraphPad Prism 10 (GraphPad Software) with a priori significance level (α) set at 0.05. We performed Shapiro-Wilk’s test for normality and Levene’s test for equal variance of the data. Statistical analyses and the number of animals per group for each experiment are detailed in the respective figure legends. Data are presented as means ± SEM. Comparisons were made using one-way or two-way (genotype × surgery) ANOVA, followed by Dunnett’s, Sidak’s, or Tukey’s post hoc tests, as appropriate.

## Declarations

### Ethics Approval

All animal studies were conducted in accordance with Washington University in St. Louis Institutional Animal Care and Use Committee.

### Consent for Publication

Not applicable.

### Availability of Data and Materials

The datasets generated and/or analyzed during the current study are not publicly available due to confidentiality restrictions related to unpublished findings and institutional policies. However, the data are available from the corresponding author upon reasonable request.

### Competing interests

FG is an employee and founder of Cytex Therapeutics, Inc.

## Funding

This work was supported by the Shriners Hospitals for Children and the National Institutes of Health (F31 AR079260, AG15768, AG46927, AR080902, AR072999, P30 AR073752, P30 AR074992, R00 AR078949).

### Author contributions

EE, KC, and FG designed the research study. EE, KC, KL, LB performed the research. EE, KL, SP, SA, AB, SD, and KC analyzed the data. EE wrote the first draft of the paper. All authors contributed to editorial changes in the manuscript. All authors read and approved the final manuscript. All authors have participated in the work and agreed to be accountable for all aspects of the work.

#### Abbreviations

OA: Osteoarthritis
PTOA: Post-traumatic osteoarthritis
DMM: Destabilization of the Medial Meniscus cKO: Conditional knockout
WT: Wildtype
SMALGO: Small animal algometer EVF: Electronic on Frey
MGS: Mouse grimace scale
μCT: Micro-computed tomography
BV/TV: Bone volume/total volume
BS/BV: Bone surface/bone volume
Tb.Th: Trabecular thickness
Tb.N: Trabecular number
Tb.Sp: Trabecular separation
BMD: Bone mineral density
Ca^2+^: Calcium ion
ACLT: Anterior cruciate ligament transection
Acan: Aggrecan

## Acknowledgments

The authors would like to thank Sara Oswald for her invaluable laboratory support. We also acknowledge the WashU Musculoskeletal Research Center Histology Core for their assistance in processing samples and Core B for providing instruments for pain testing

