## Supplementary Figures for "Chondrocyte-Specific Knockout of *Piezo1* and *Piezo2* Protects Against Post-Traumatic Osteoarthritis Structural Damage and Pain in Mice"

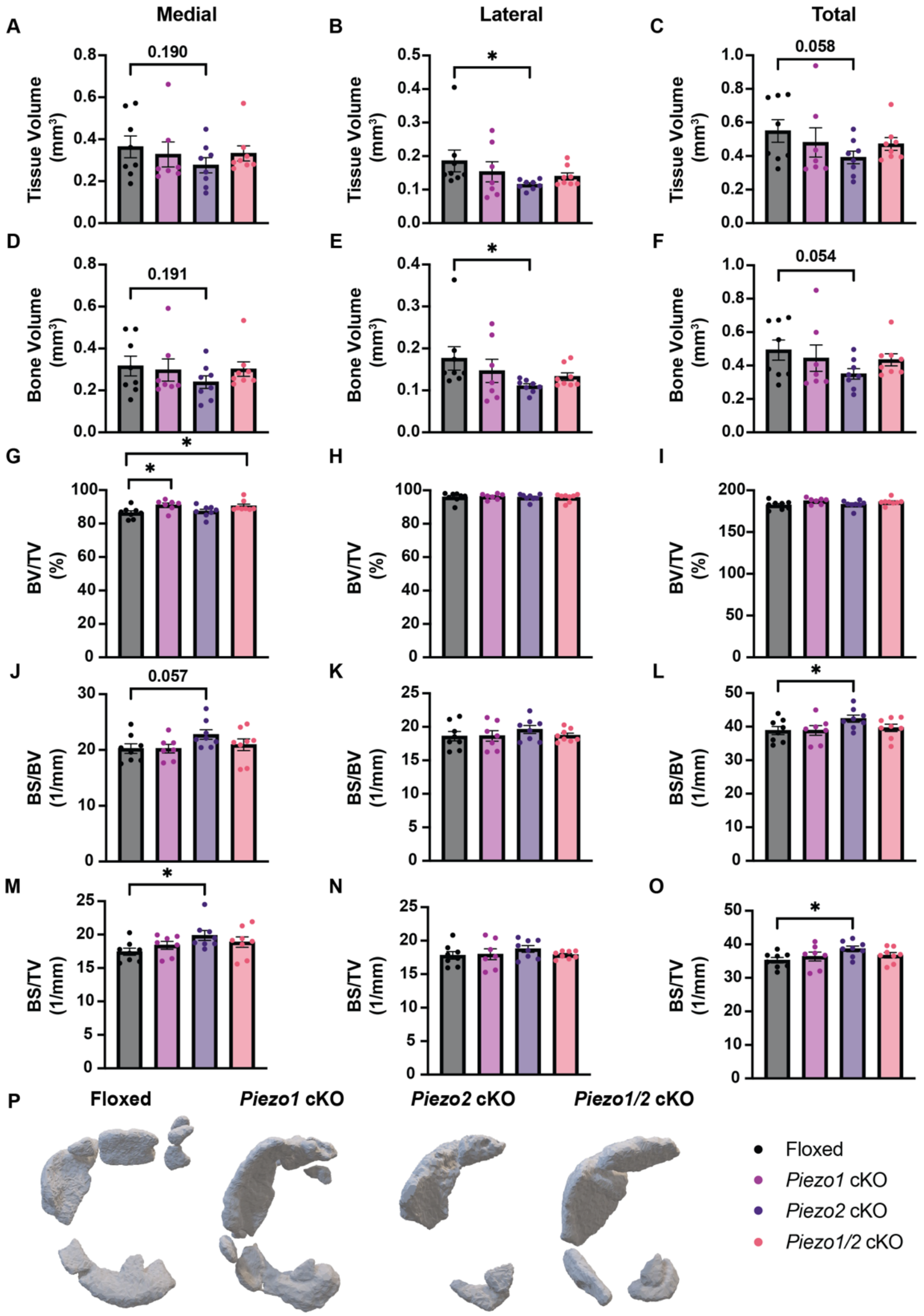


**Supplemental Figure 1. Meniscus Ossification Quantification.** (A-C) Tissue volume (mm³), (D-F) Bone volume (mm³), (G-I) Bone volume fraction (BV/TV, %), (J-L) Bone-surface-to-volume ratio (BS/BV) (1/mm), and (M-O) Bone surface density (BS/TV) (1/mm). are shown for medial (A, D, G, J, M), lateral (B, E, H, K, N), and total (C, F, I, L, O) meniscus. Black bars represent Floxed control mice, pink bars represent *Piezo1* cKO mice, purple bars represent *Piezo2* cKO mice, and orange bars represent *Piezo1/2* cKO mice. Statistical comparisons were performed using an unpaired t-test. Data are presented as mean ± SEM. (P) Representative 3D models of medial meniscus from Floxed, *Piezo1* cKO, *Piezo2* cKO and *Piezo1/2* cKO animals.

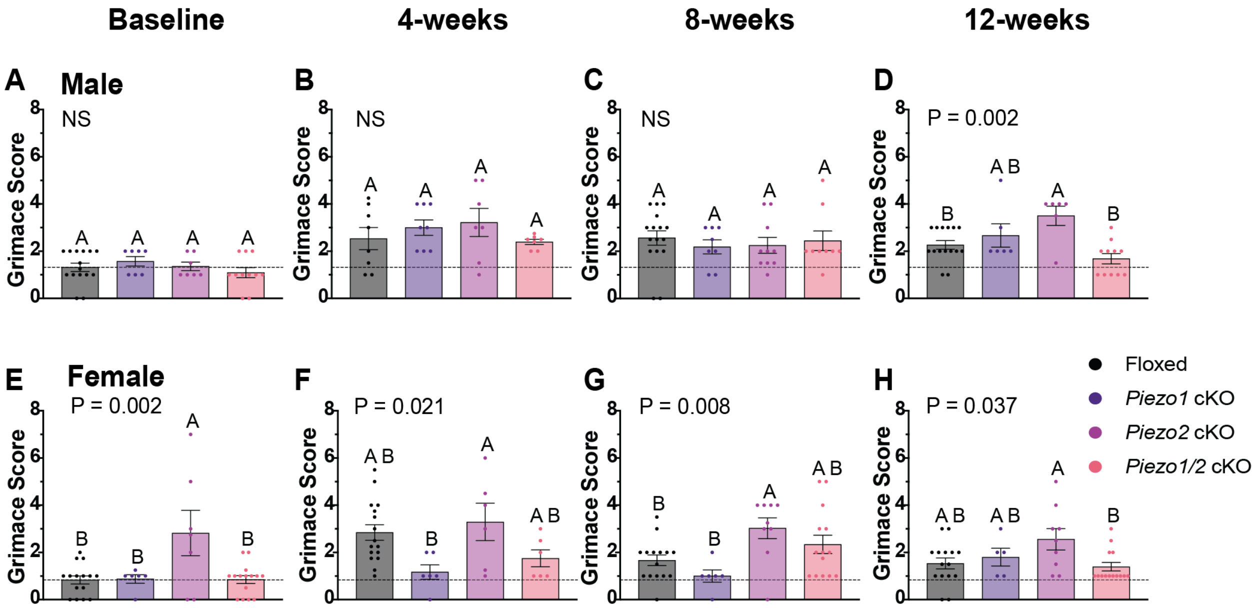


**Supplemental Figure 2. Mouse Grimace Scale (MGS) Assessment of Pain in Male and Female Mice Post-DMM Surgery**

Pain-related behaviors in male and female Floxed, *Piezo1* cKO (P1 cKO - purple), *Piezo2* cKO (P2 cKO - pink), and *Piezo1/2* cKO (P1/2 cKO - orange) mice were assessed using the Mouse Grimace Scale (MGS) at baseline (0 weeks), and at 4, 8, and 12 weeks post-DMM surgery. Panels A-D show the results for male mice, while panels E-H show the results for female mice. A one-way ANOVA was used for statistical analysis, followed by Tukey's multiple comparisons test to assess differences between genotypes. At baseline, female *Piezo2* cKO mice had significantly higher grimace scores compared to other genotypes ($P = 0.002$), indicating increased pain sensitivity (E). At 12 weeks post-DMM, *Piezo2* cKO male mice exhibited higher pain scores compared to Floxed and *Piezo1/2* cKO groups ($P = 0.002$) (D), while female *Piezo1/2* cKO mice had significantly lower pain scores compared to *Piezo2* cKO mice ($P = 0.037$) (H). Across all timepoints, *Piezo1/2* cKO mice generally showed reduced pain levels compared to *Piezo2* cKO, suggesting a potential protective effect in double knockout mice. Different letters above bars indicate significant differences between groups ($P < 0.05$). Error bars represent mean ± SEM. The dashed line represents baseline grimace scores.

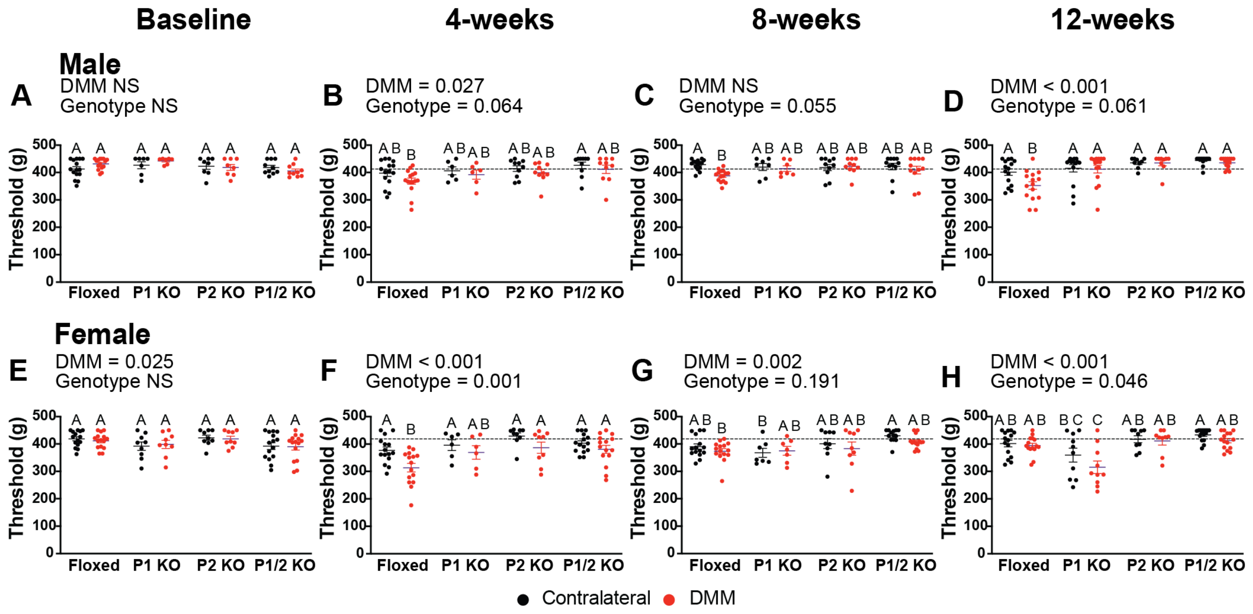


**Supplemental Figure 3. Pressure-Pain Hyperalgesia Thresholds in Male and Female Mice Post-DMM Surgery.** Pressure-pain hyperalgesia thresholds were evaluated using the SMALGO in both DMM (red) and contralateral limbs (black) of male (A-D) and female (E-H) Floxed (black), *Piezo1* cKO (P1 cKO - purple), *Piezo2* cKO (P2 cKO – pink), and *Piezo1/2* cKO (P1/2 cKO – orange) mice at baseline, 4, 8, and 12 weeks post-DMM. A two-way ANOVA was used for statistical analysis (n=6-15/genotype/sex). At baseline, no significant differences were observed among genotypes. At 12 weeks post-DMM, all *Piezo c*KO male groups have significantly higher DMM thresholds compared to Floxed mice. In female mice, a *Piezo2 c*KO results in decreased thresholds compared to all groups. Different letters indicate statistically significant differences between groups (P < 0.05). Error bars represent mean ± SEM. The dashed line represents baseline pressure-pain hyperalgesia.

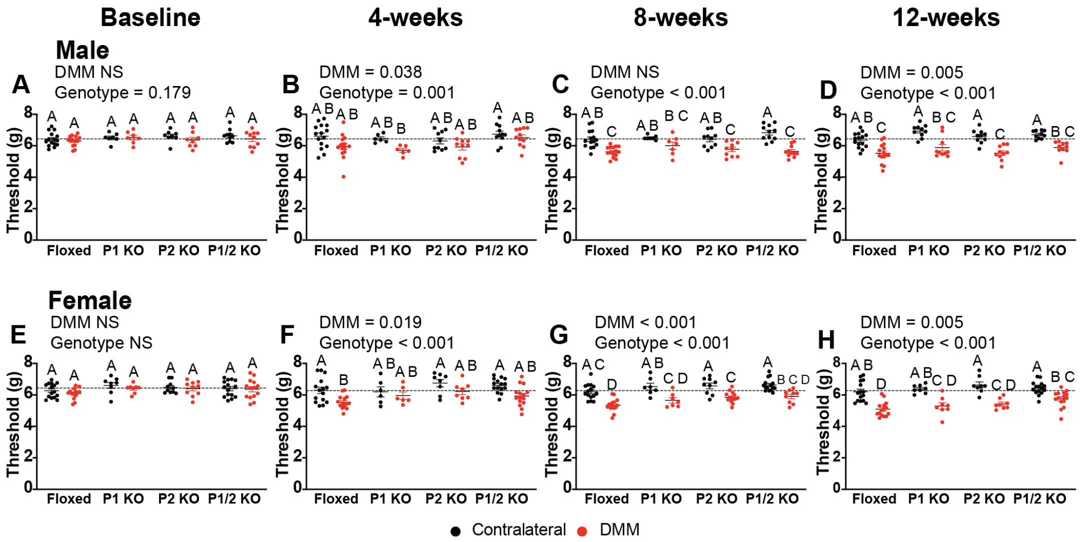


**Supplemental Figure 4. Tactile Allodynia Thresholds in Male and Female Mice Post-DMM Surgery.** Tactile allodynia thresholds were evaluated using the Electronic Von Frey (EVF) in both DMM (red) and contralateral limbs (black) of male (A-D) and female (E-H) Floxed, *Piezo1* cKO (P1 KO), *Piezo2* cKO (P2 KO), and *Piezo1/2* cKO (P1/2 KO) mice at baseline, 4, 8, and 12 weeks post-DMM. A two-way ANOVA was used for statistical analysis (n=6-15/genotype/sex). At baseline, no significant differences were observed among genotypes. At 12 weeks post-DMM, all male groups had significantly lower DMM thresholds compared to contralateral limbs, with no significant differences across genotypes. In female mice, *Piezo1/2* cKO DMM thresholds were significantly higher compared to Floxed mice, indicating a protective effect. Error bars represent mean ± SEM. Different letters indicate statistically significant differences between groups ($P < 0.05$). The dashed line represents baseline EVF thresholds.

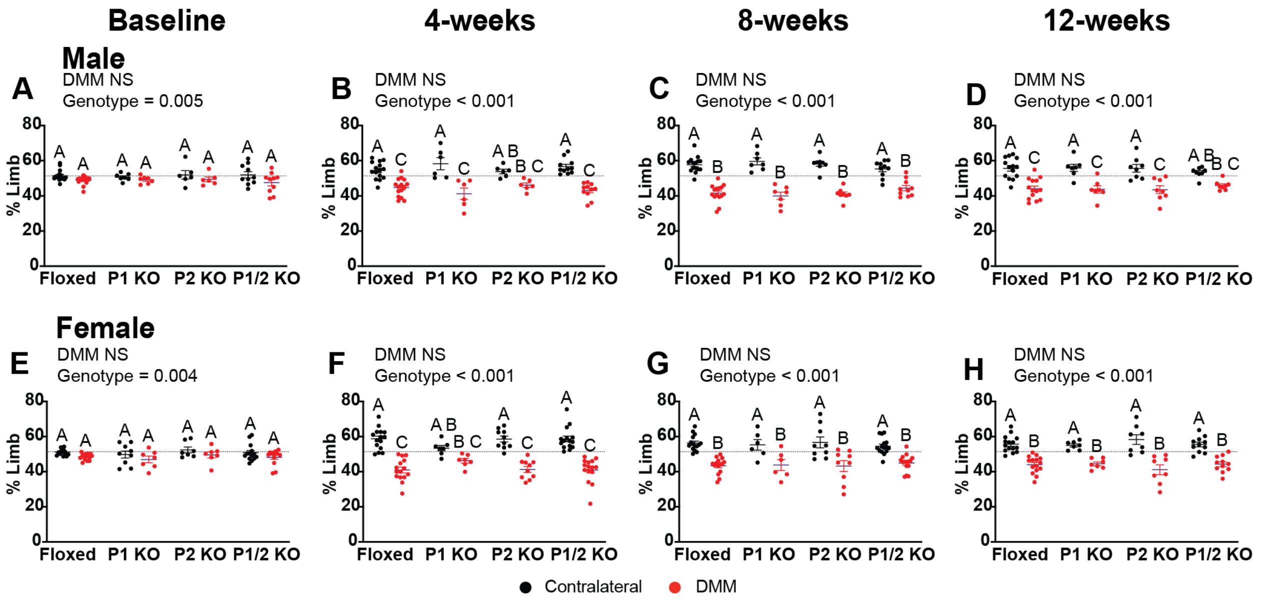


**Supplemental Figure 5. Static Weight Bearing in Male and Female Mice Post-DMM Surgery.** Static weight bearing measurements were evaluated in both DMM (red) and contralateral limbs (black) of male (A-D) and female (E-H) Floxed (black), *Piezo1* cKO (P1 KO), *Piezo2* cKO (P2 KO), and *Piezo1/2* cKO (P1/2 KO) mice at baseline, 4, 8, and 12 weeks post-DMM. A two-way ANOVA was used for statistical analysis (n=6-15/genotype/sex). At baseline, a significant effect of genotype was observed in both male (P = 0.005) and female (P = 0.004) mice. At 4 weeks post-DMM, static weight bearing was significantly impaired in all groups, with *Piezo2* cKO males and *Piezo1* cKO females showing no significant differences between limbs. At 8 weeks, all genotypes demonstrated a significant reduction in weight bearing in the DMM limb, suggesting impaired load distribution following DMM surgery. By 12 weeks, differences in weight distribution remained significant across most genotypes, except for *Piezo1/2 c*KO male mice, highlighting the persistent effects of DMM surgery. Error bars represent mean ± SEM. Different letters indicate statistically significant differences between groups (P < 0.05). The dashed line represents baseline load distribution.

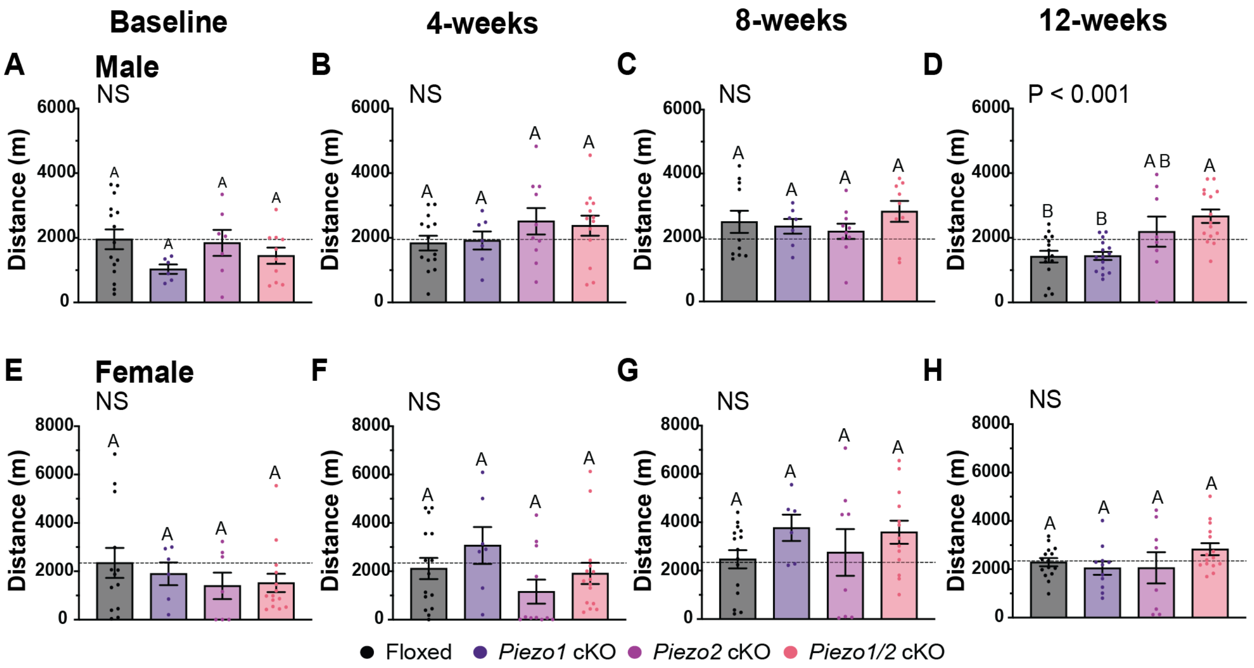


**Supplemental Figure 6. Voluntary Wheel Running in Male and Female Mice Following DMM Surgery.** Voluntary wheel running distances (in meters) were assessed at baseline (A, E), 4 weeks (B, F), 8 weeks (C, G), and 12 weeks (D, H) post-DMM surgery in four groups: Floxed (black), *Piezo1* cKO (P1 cKO - purple), *Piezo2* cKO (P2 cKO - pink), and *Piezo1/2* cKO (P1/2 cKO - orange). Male mice are shown in (A-D), and female mice in (E-H). A significant difference in distance run was detected among male groups at 12 weeks ($P < 0.001$) (D), where *Piezo1/2* cKO mice exhibited significantly greater activity compared to *Piezo1* cKO and Floxed controls. For female mice, no significant differences were observed across genotypes at any timepoint, with all groups demonstrating comparable recovery in running distances post-DMM surgery. The dashed line indicates baseline control distance for male and female mice. Different letters indicate statistical significance, one-way ANOVA with Tukey's multiple comparisons test). Error bars represent mean ± SEM.

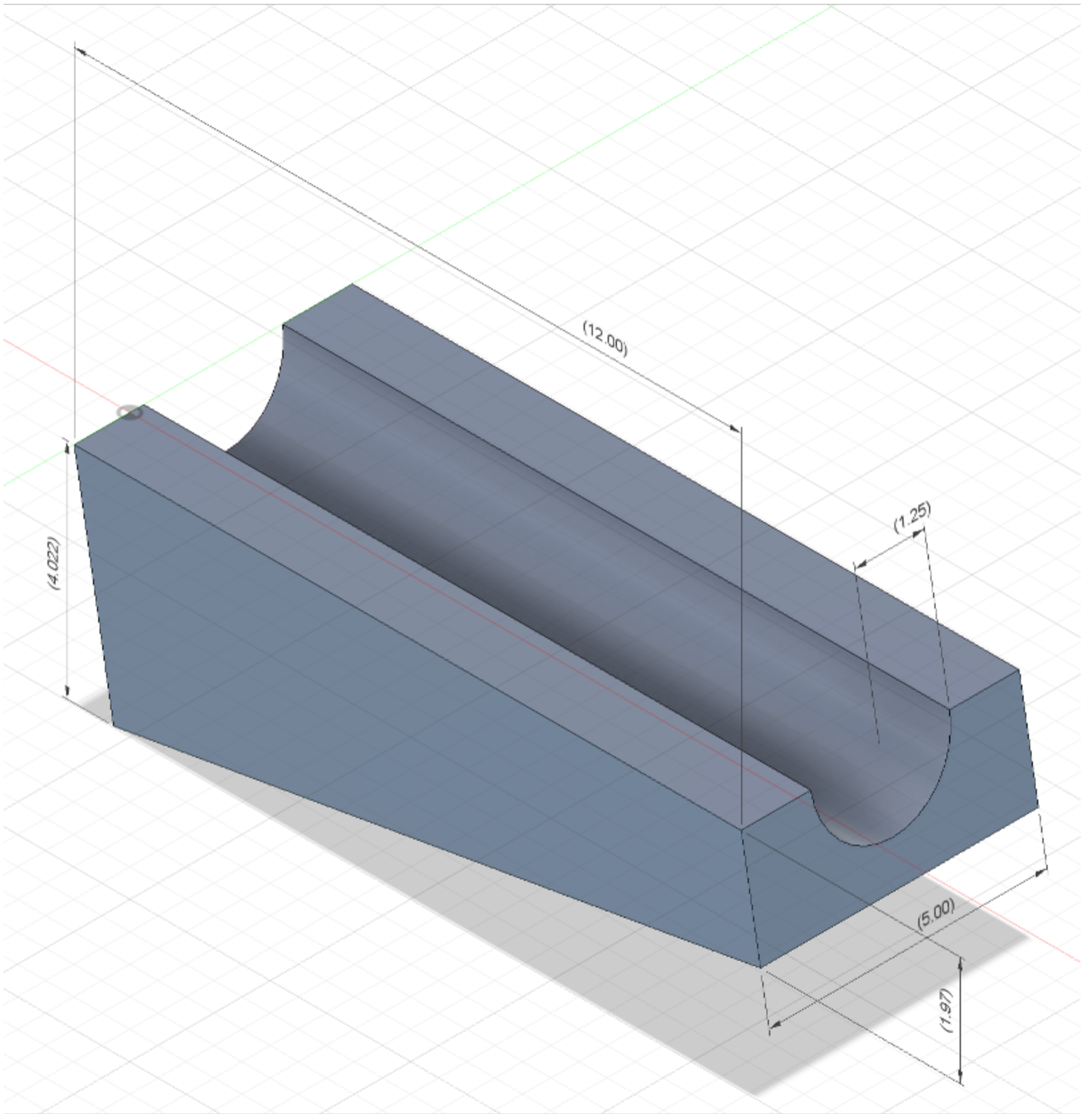


**Supplemental Figure 7. Custom Rig for Calcium Imaging.** CAD rendering of a custom imaging rig designed to stabilize femoral condyles at a 10° angle. The rig features a contoured groove to securely position the femur, ensuring consistent orientation for imaging. Dimensions are provided in millimeters.

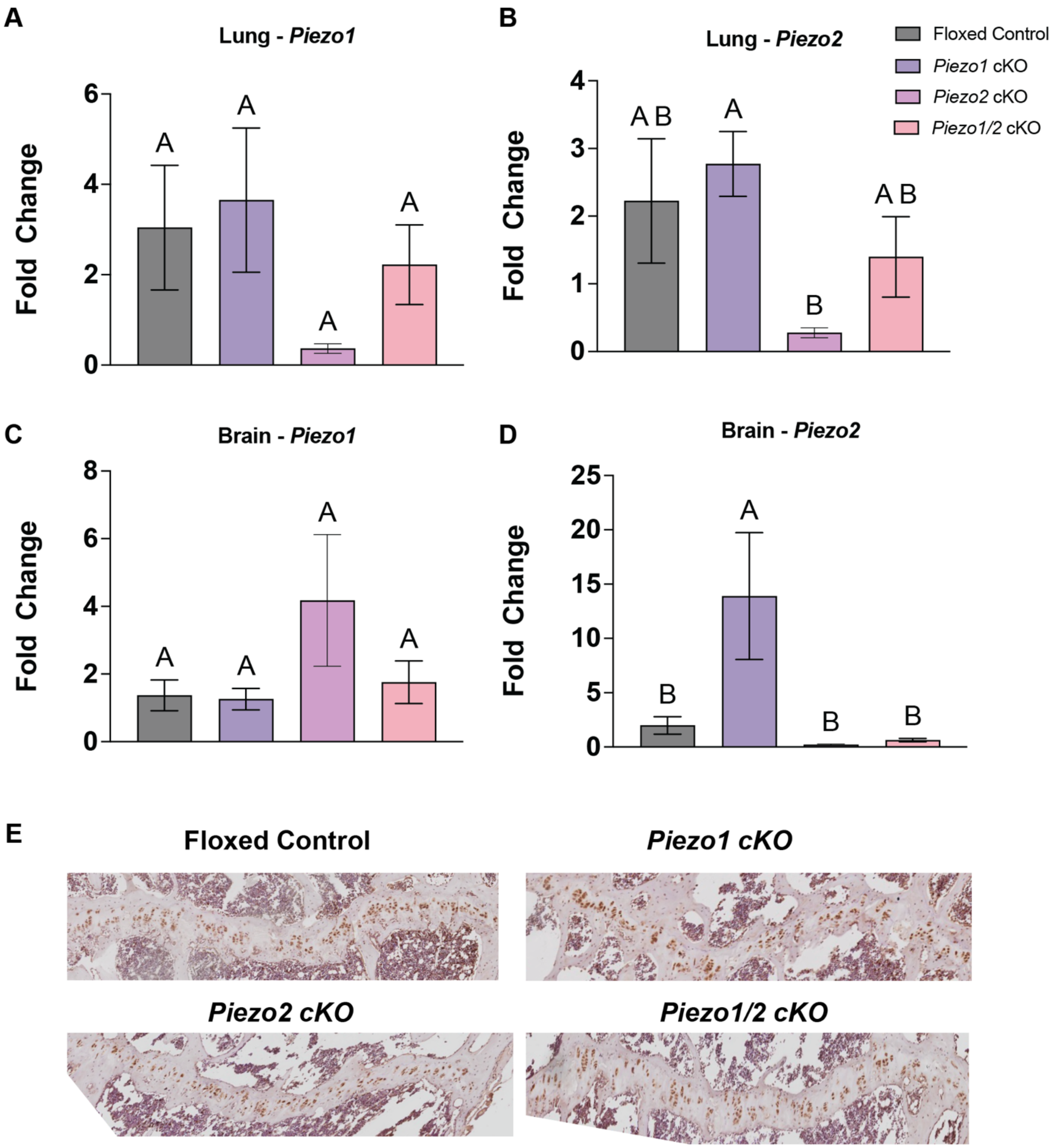


**Supplemental Figure 8. Piezo Expression in Non-Cartilage Tissues.** Quantitative PCR (qPCR) was performed on murine lung (A,B) and brain (C,D) tissues from floxed (black), *Piezo1* cKO (purple), *Piezo2* cKO (pink), and *Piezo1/2* cKO (orange) mice at 28 weeks of age (n = 6–10/genotype). *Piezo1* expression was measured in A and C and *Piezo2* expression was measured in B and D. Data are represented as fold change. Data analyzed using 1-way ANOVA. Statistical differences are represented by different letters. Femoral growth plate samples were stained using Immunohistochemistry for PIEZO2. Positive protein expression is seen through brown DAB expression.
